# Thymine DNA Glycosylase Binds to R-Loops and Excises 5-Formyl and 5-Carboxyl Cytosine from DNA/RNA Hybrids

**DOI:** 10.1101/2025.08.05.668694

**Authors:** Baiyu Zhu, Lakshmi S. Pidugu, Mary E. Cook, Xinyu Y. Nie, E. A. P. Tharaka Amarasekara, Jerome S. Menet, Alexander C. Drohat, Jonathan T. Sczepanski

## Abstract

R-loops are three-stranded nucleic acid structures consisting of a DNA/RNA hybrid and a displaced single-stranded DNA. Once considered rare byproducts of transcription, R-loops are now recognized as important regulators of various nuclear processes. In particular, evidence indicates a role for R-loops in regulating DNA methylation dynamics. R-loops have been shown to promote active DNA demethylation—the enzymatic reversal of 5-methylcytosine (5mC) back into cytosine—by recruiting associated proteins, providing an attractive targeting mechanism. Nevertheless, many important aspects of this process, including whether the associated proteins bind to and function on R-loops, remain to be substantiated. In this study, we demonstrate for the first time that thymine DNA glycosylase (TDG), a key enzyme in the active DNA demethylation pathway, binds tightly to R-loops in vitro and can excise DNA demethylation intermediates 5-formylcytosine (5fC) and 5-carboxycytosine (5caC) from DNA in DNA/RNA hybrid duplexes. We also show that R-loops guide the strand-specific activity of TDG at CpG sites, potentially explaining the asymmetric distribution of 5fC/5caC at gene promoters. Furthermore, we provide important mechanistic insights into base excision on DNA/RNA hybrid duplexes using ^19^F NMR. Finally, our findings suggest that TDG–R-loop interactions may be widespread in human cells. Collectively, our results provide strong evidence that R-loops play a critical role in DNA demethylation and support a mechanism in which 5fC/5caC are directly removed from DNA/RNA hybrids in cells.

**GRAPHICAL ABSTRACT:** 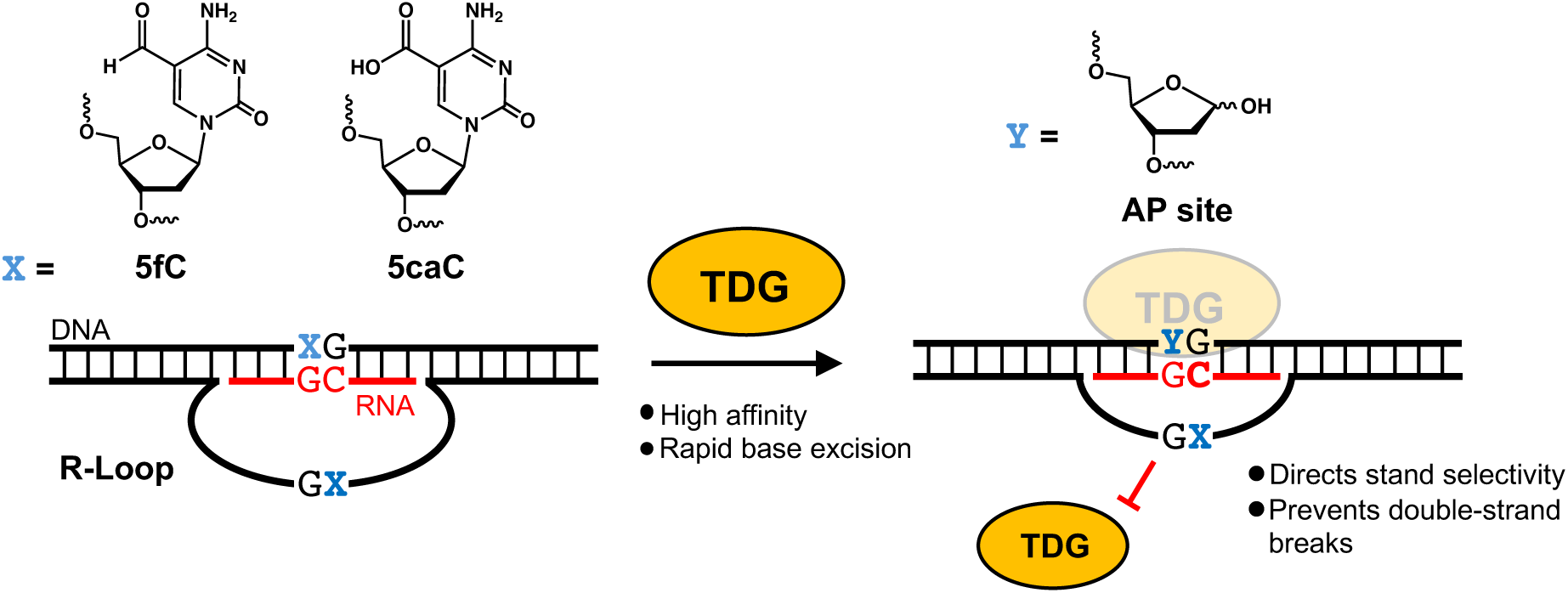

## INTRODUCTION

5-methylcytosine (5mC) is a highly conserved epigenetic modification of DNA found predominantly within CpG dinucleotides and is associated with heterochromatin formation leading to silencing of gene expression. While traditionally viewed as a static DNA modification, recent research has revealed that 5mC is dynamic in nature, and reversal of 5mC back into cytosine (i.e. demethylation) is an essential biological pathway in mammals. The reversal process involves two primary mechanisms: passive and active DNA demethylation. Passive DNA demethylation occurs when the methylated DNA is progressively diluted through successive rounds of replication, primarily due to the deactivation or nuclear exclusion of maintenance DNMT1 or its co-factors. Alternatively, the enzymatic reversal of 5mC to cytosine (referred to as “active” DNA demethylation) involves the successive oxidation of 5mC into 5-hydroxymethylcytosine (5hmC), 5-formylcytosine (5fC), and 5-carboxycytosine (5caC) by the ten-eleven translocation (TET) family of dioxygenases (Figure 1).(1–5) Thymine DNA glycosylase (TDG) excises the two most oxidized cytosine derivatives, 5fC and 5caC, from DNA (4,6), thereby initiating base excision repair (BER) and ultimately restoring unmodified cytosine.(7–14) The active DNA demethylation pathway is essential for maintaining proper epigenetic states during development, regulating hormone-dependent gene expression in differentiated cells, and preventing tumor formation. Therefore, a comprehensive understanding of the active DNA demethylation pathway and its component enzymes is crucial for deciphering how DNA methylation landscapes are established in both normal and disease conditions.

**Figure 1.**
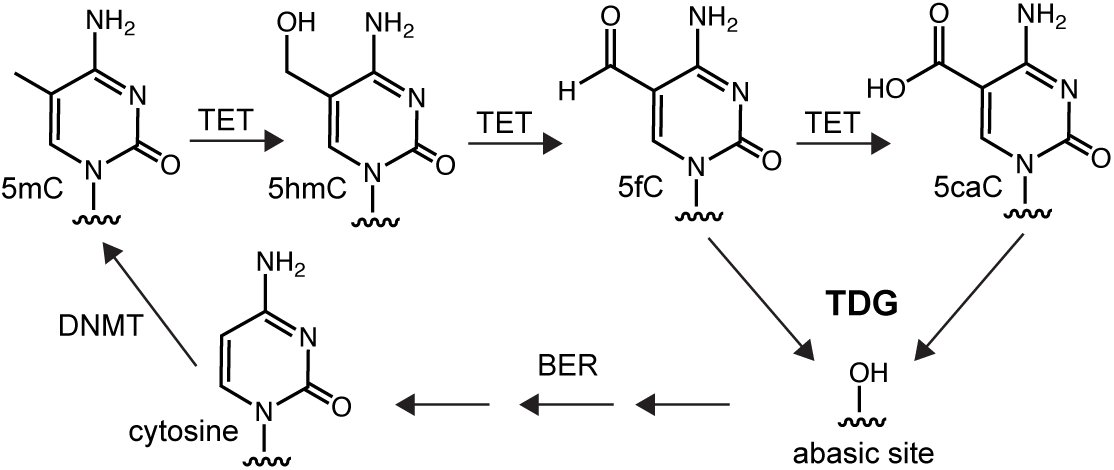
The active DNA demethylation mediated by TET and TDG.

TDG is a member of the uracil DNA glycosylase (UDG) superfamily that was originally discovered to excise pyrimidine bases from G•U and G•T mispairs and to initiate BER at these sites.(15,16) This activity is thought to help protect the genome against C→T transition mutations that arise via the deamination of cytosine and 5-methylcytosine, respectively. Importantly, as the only mammalian glycosylase currently known to be capable of removing 5fC and 5caC from DNA, TDG plays a central role in the active DNA demethylation pathway.(3,17) Depletion of TDG from mouse embryonic stem cells (mESCs) results in the accumulation of 5fC and 5caC at regulatory sites, such as promoters and enhancers, pointing to a role for TDG in epigenetic regulation of gene expression.(13,14,18–21) TDG’s catalytic activity is essential during development for preserving the epigenetic integrity of numerous developmental and tissue-specific genes; its loss in the germline results in embryonic lethality.(11,12) Beyond development, TDG-mediated DNA demethylation plays a key role during transcriptional activation of a subset of hormone-responsive genes in differentiated cells.(22–26) Importantly, evidence suggests that 5fC and 5caC function as stable epigenetic modifications to regulate gene expression in addition to their roles as intermediates in the DNA demethylation pathway.(27–32) Thus, TDG has a pivotal role of determining whether 5fC and 5caC are retained as potential epigenetic marks or removed via BER.

A key bottleneck in our current understanding of the active DNA demethylation pathway is that it remains unclear how the associated enzymes, including TDG, are targeted and regulated at the genome level. DNA demethylation occurs at specific promoters and enhancers in response to developmental or environmental cues, typically affecting only a limited number of CpG dinucleotides.(11,12,33) Yet, it is unknown how this precise targeting is achieved. While most studies aimed at answering this question have focused on the interactions of the demethylation machinery with its protein partners, such as sequence-specific transcription factors, emerging evidence now indicates potential regulatory relationships with RNA.(26,34,35) Indeed, several proteins involved in controlling DNA methylation dynamics, including DNA methyltransferases (DNMT1, DNMT3a/b), TETs, TDG, and GADD45A (growth arrest and DNA-damage-inducible, alpha)(25,36–39) are localized at target sites in an RNA-dependent manner. One possible mechanism for these regulatory functions involves the formation of R-loops. R-loops are three-stranded nucleic acid structures consisting of a DNA/RNA hybrid and a displaced single-stranded DNA (ssDNA). While prevalent during transcription, R-loops have gained recent attention for having important roles in many other nuclear processes, including recombination and DNA repair. Importantly, evidence indicates a role for R-loops in regulating DNA methylation dynamics. Genome-wide mapping studies show that the formation of R-loops negatively correlates with DNA methylation, especially at gene promoters (40), consistent with their ability to inhibit the binding and catalytic activity of DNA methyltransferases (DNMTs). Other evidence suggests that R-loops can promote 5mC reversal. Notably, Arab et al. showed that several long noncoding RNAs (lncRNAs) form R-loops near transcriptional start sites (TSS) and demonstrated that these structures can promote local DNA demethylation and gene expression.(41) The proposed mechanism involves specific recognition of R-loops by the scaffolding protein GADD45A (growth arrest and DNA-damage-inducible, alpha), which in turn recruits the DNA demethylation machinery, including TET1 and TDG, to the underlying CpG sites for processing. Because R-loops are formed in a sequence-specific manner, this provides an attractive mechanism for the precise targeting of CpGs for demethylation. Interestingly, many of the R-loops implicated in directing DNA demethylation activities overlap with target CpG sites, suggesting that DNA demethylation may occur directly on these unique hybrid structures. In the case of TDG, for example, this would require recognition and base excision of 5fC and 5caC on DNA/RNA hybrid substrates. However, whether TDG or other components of the demethylation machinery function on R-loops remains to be substantiated.

To address these questions, we investigated the activity of TDG on R-loop substrates for the first time. We show that TDG binds R-loops in vitro and is capable of excising 5fC and 5caC from DNA/RNA hybrid duplexes, supporting a mechanism whereby DNA demethylation occurs directly over R-loops. Furthermore, we demonstrate that R-loops can direct the strand selectivity of TDG and protect against double-strand break formation at symmetrically modified CpG dinucleotides. Finally, we provide evidence showing that interactions between R-loops and TDG are extensive in human cells. Together, our results advance the understanding of the role R-loops play in epigenetic regulation, particularly in the context of TDG-mediated DNA demethylation.

## MATERIALS AND METHODS

### Reagents

All synthetic oligonucleotides were either purchased from Integrated DNA Technologies (Coralville, IA, USA) or prepared by solid-phase synthesis on an Expedite 8909 DNA/RNA synthesizer. Nucleoside phosphoramidites and DNA synthesis reagents were purchased from Glen Research (Sterling, VA, USA). Sulfo-Cyanine3 (Cy3; cat. No: 21320) and Cyanine5 (Cy5; cat. No: 23320) NHS ester dyes were purchased from Lumiprobe Life Science Solutions (Hunt Valley, MD, USA). PIPES, 1M, pH 8.0 (cat. no: J60618.AK), Sodium Deoxycholate Detergent (cat. no: 89904), N-Lauroylsarcosine sodium salt, 95% (cat. no: J60040.09), Dynabeads Protein A for Immunoprecipitations (cat. no: 10001D), NP-40 Surfact-Amps Detergent (cat. no: 85124), Halt Protease Inhibitor cocktail (cat. no: 78430), NuPAGE LDS Sample Buffer (cat. no: NP0007), Goat anti-Rabbit IgG (H+L) Highly Cross-Adsorbed Secondary Antibody, Alexa Fluor Plus 647 (cat. no: A32733), Lipofectamine 3000 Transfection Reagent (cat. no: L3000008) were purchased from ThermoFisher Scientific (Waltham, MA, USA). Hpy188I (cat. no: R0617S) and proteinase K (cat. no: P8107S) were purchased from New England Biolabs (Ipswich, MA, USA). Mouse IgG2a Isotype Control (cat. no: M5409-.1MG) and BL21Rosetta (DE3) (cat. no: 70954-3) were purchased from Sigma Aldrich (Rockville, MD, USA). Anti-DNA-RNA Hybrid [S9.6] Antibody (cat. no: ENH001) was purchased from Kerafast, Inc. (Newark, CA, USA). TDG Polyclonal antibody (cat. no: 13370-1-AP) was purchased from Proteintech (Rosemont, IL, USA). Dry Powder Milk (cat. no: M17200-500.0) was purchased from RPI-Research Products International (Mount prospect, IL, USA).

### Biological Resources

HeLa S3 cells (cat. no: CCL-2.2) were purchased from American Type Culture Collection (Manassas, VA) and cultured in Dulbecco’s Modified Eagle’s Medium (DMEM; Thermo Fisher Scientific, Waltham, MA) supplemented with 25 mM HEPES, 1 mM GlutaMax, 10% FBS, and 100U/mL Penicilin-Streptomycin (Thermo Fisher Scientific, Waltham, MA). Cells were maintained at 37 °C in humidified CO_2_ (5%) atmosphere. The TDG mammalian expression vector (cat. no: HG13000-UT) was purchased from Sino Biological, Inc. (Wayne, PA, USA).

### Protein expression and purification

Full-length human TDG was expressed and purified as described previously with minor modifications.(42) In brief, the TDG plasmid (pET28a-hTDG; 35 ng) was transformed into BL21 Rosetta (DE3) cells and the outgrowth was used to prepared 4 × 25 mL cultures of Luria-Bertani broth (LB) supplemented with 50 ug/mL Kanamycin and Chloramphenicol. Following overnight shaking at 37 °C, 10 mL of the overnight culture was added to 1 L of LB supplemented with 50 ug/mL Kanamycin and chloramphenicol. The cells were induced with 0.25 mM IPTG at 15 °C for 15 hours when reached OD∼0.6. They were then pelleted by centrifugation at 4500 rpm for 20 minutes at 4 °C using Sorvall RC-5C plus centrifuge. The resulting pellets were stored at −80 °C overnight and then thawed at 4 °C. The cell pellets were then resuspended in lysis buffer (50 mM PO_4_^3-^ pH 8, 300 mM NaCl, 5 mM Imidazole, 1 mM BME) supplemented with protease inhibitor. To start lysis, lysozyme (1 mg/mL) and DNase (0.025 U/uL) were added to the resuspended cell pellets and kept on ice for 30 minutes. The cells were sonicated for 6 minutes. The resulting lysates were centrifuged (10,000 RPM, 60 minutes) and filtered with a 0.2 um syringe tip filter (cat. no: SLGPR33RS, Millipore Sigma, Burlington, VT, USA). The pre-packed His GraviTrap TALON column (cat. no: 29000594, CYTIVA, Wilmington, DE, USA) was equilibrated with 25 mL of lysis buffer twice and then the filtered lysate was passed through. The TDG bound resin was then washed with 30 mL of wash 1 (700 mM NaCl in lysis buffer) and again with 30 mL of lysis buffer. TDG was eluted with 3 x 5 mL of elution buffer (500 mM Imidazole in lysis buffer). The eluted TDG was exchanged into IEA buffer (20 mM HEPES pH 7.5, 75 mM NaCl, 1 mM DTT, 0.2 mM EDTA, 1% Glycerol) using a 5 mL Sephadex G-25 resin HiTrap Desalting column (cat. no: 17140801, CYTIVA, Wilmington, DE, USA). TDG was loaded onto a 1 mL HiTrap Q HP anion exchange chromatography column (cat. no: 17115301, CYTIVA, Wilmington, DE, USA) that was equilibrated with IEA buffer. Bound TDG was then eluted using a linear gradient (0-100%) of IEB buffer (20 mM HEPES pH 7.5, 1 M NaCl, 1 mM DTT, 0.2 mM EDTA, 1% Glycerol) in 500 uL fractions. Fractions containing TDG were pooled and concentrated using an Amicon Ultra centrifugal filter with a 3 kDa MWCO (cat. no: UFC900308, Millipore Sigma, Burlington, VT, USA).

TDG^82-308^ was expressed in *E. coli* BL21(DE3) at (22 °C) and purified (at 4 °C) by Ni-affinity, ion-exchange (SP sepharose) and size exclusion chromatography as described previously.(43,44) Enzyme purity was >99% as judged by SDS-PAGE with Coomassie staining, and the concentration was determined by absorbance (280 nm) using an extinction coefficient of ε^280^ = 17.4 mM^−1^cm^−1^.(45,46) Purified enzyme was flash frozen and stored at −80 °C.

### Oligonucleotide synthesis and purification

All oligonucleotides used in this study are shown in Table S1. Synthetic oligonucleotides prepared in-house were made using an Expedite 8909 DNA/RNA synthesizer according to manufacturer’s recommended protocol. All oligonucleotides (purchased or synthesized in-house) were purified by denaturing polyacrylamide gel electrophoresis (PAGE) (20%, 19:1 acrylamide:bisacrylamide). Targets bands were excised from the gel and eluted overnight at room temperature in elution buffer (200 mM NaCl, 10 mM EDTA, 10 mM Tris-HCl pH 7.6). Samples were then filtered to remove gel fragments and desalted by ethanol precipitation. 6-carboxyfluorescein (FAM)-labeled oligonucleotides were either purchased directly or synthesized in-house using the 5’-fluorescein phosphoramidite (cat. no: 10-5901-90E, Glen Research, Sterling, VA, USA). Sulfo-Cy3 and Sulfo-Cy5 labeling was carried out using the corresponding NHS esters. Oligonucleotides harboring a 5′-amino modifier were purchased from Integrated DNA Technologies (Coralville, IA, USA). Labelling reactions were performed by mixing the amine-modified oligonucleotide with 10-fold molar excess of the indicated NHS ester dye in 0.1 M sodium bicarbonate buffer (pH 8.5). The reaction was kept on ice overnight and the excess dye was subsequently removed by ethanol precipitation prior to use. Purified oligonucleotides were dissolved in water and their concentrations were determined by absorbance at 260 nm using a NanoDrop 2000c (ThermoFisher Scientific, Waltham, MA). Duplex and R-loop substrates were prepared by annealing 10 uM each of the corresponding strands (Table S1) in annealing buffer (50 mM NaCl, 10mM Tris-HCl pH 7.5). The mixture was kept at 95°C for 3 minutes before being cooled down to room temperature over the course of 90 minutes.

### Electrophoretic mobility shift assay (EMSA)

EMSAs were carried out as described previously with minor modifications.(42) Briefly, the indicated substrate (5 nM) was mixed with increasing concentrations of TDG (0 – 300 nM) in binding buffer (100 mM NaCl, 2.5 mM MgCl_2_, 10 mM Tris-HCl pH 7.5, and 5% glycerol). The reaction mixture was incubated at 30 °C for 30 minutes and an aliquot was resolved by 0.6% agarose gel buffered with 1 × TBE. Electrophoresis was carried out for 1 hour (6–8 V/cm) at 4 °C. The gel was visualized using a ChemiDoc^TM^ MP Imaging System (Bio-Rad Laboratories, Inc., Hercules, CA, USA) and images were quantified using Image Lab software version 6.1.0 (Bio-Rad Laboratories, Inc.). GraphPad Prism 9 Version 9.4.1 was used to fit equations for specific binding with Hill slope.

### Glycosylase assays

Single-turnover kinetic reactions were initiated by mixing 100 nM of the indicated substrate with 1 uM of TDG in a buffer containing 100 mM NaCl, 2.5 mM MgCl_2_, and 10 mM Tris-HCl (pH 7.5). Aliquots (2 uL) were removed at the desired time points and added to a solution (2 uL) of 1% SDS in water to quench the reaction. To cleave the abasic site product, equal volume of 0.2 M NaOH was added to the aliquots, which were subsequently heated at 70 °C for 5 minutes before adding 8 uL of denaturing loading buffer (90% formamide, 10 mM EDTA pH 8). The products were then resolved using denaturing PAGE (20%, 19:1 acrylamide:bisacrylamide). The gels were visualized by ChemiDoc^TM^ MP Imaging System (Bio-Rad Laboratories, Inc., Hercules, CA) and images were quantified using Image Lab version 6.1.0 (Bio-Rad Laboratories, Inc., Hercules, CA). The fraction product versus time were fitted to eq. 1:

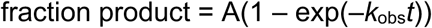

Where A is the amplitude, *k*_obs_ is the rate constant, and *t* is the reaction time. Experiments were performed with saturating TDG, which was confirmed by obtaining similar rate constants for experiments carried out at higher enzyme concentrations.

### ^19^F NMR spectroscopy

The ^19^F NMR experiments were performed at 25 °C on a Bruker 600 MHz spectrometer (564.2 MHz for ^19^F) equipped with four channels, a Z-axis gradient, and a 5 mm HFCN cryogenic probe (optimized for ^1^H, ^19^F, ^13^C and ^15^N), as previously described.(47,48) ^19^F NMR experiments were carried out using a construct of TDG comprised of residues 82–308 (TDG^82-308^) that is functionally equivalent to that of full-length TDG. The samples contained DNA at a concentration of 62-75 μM DNA and (if present) a twofold higher concentration of TDG^82-308^. The buffer for ^19^F NMR experiments consisted of 0.1 M NaCl, 0.03 mM TCEP, 15 mM Tris-HCl pH 7.5, and 10% D_2_O. The ^19^F NMR experiments were collected with 2048 complex points, an acquisition time of 0.66 seconds, a relaxation delay of 2.0 seconds, and with 5,000 scans for free DNA and 8,000-48,000 scans for TDG complexes with DNA or DNA/RNA hybrids. The NMR data were processed by applying exponential multiplication with 25 Hz line broadening prior to Fourier transformation and baseline correction using TopSpin (Bruker) and analysed using TopSpin and CcpNmr.(49) The ^19^F chemical shift values (δ^19^F) reported herein are relative to an external sample of trifluoroacetic acid (6.5 mM) in the identical buffer.

### Double-strand break assay

Reactions were initiated by mixing 200 nM TDG and 20 nM APE1 with 100 nM of the indicated substrate in a reaction buffer containing 100 mM NaCl, 2.5 mM MgCl_2_, and 10 mM Tris-HCl (pH 7.5) at 30 °C. A control without APE1 was also setup side-by-side. Aliquots (2 uL) were removed at the indicated time points and quenched with 1.6 U of proteinase K (New England Biolabs, Ipswich, MA), followed by the addition of 50% glycerol in water (1 uL). Aliquots were left to sit at room temperature for 30 minutes. Samples were then resolved by native PAGE (10%, 29:1 acrylamid:bisacrylamide). The gels were visualized by ChemiDoc^TM^ MP Imaging System (Bio-Rad Laboratories, Inc., Hercules, CA) and quantified as described above.

### DNA-RNA immunoprecipitation (DRIP)

DRIP assays were performed with non-crosslinked HeLa S3 cells using the S9.6 antibody as previously described with minor modifications.(50) HeLa S3 cells were maintained in T-75 size CELLSTAR TC Treated Cell Culture flask (cat. no: 07-000-225, ThermoFisher Scientific, Waltham, MA, USA) in Dulbecco’s Modified Eagle’s Medium (DMEM; Thermo Fisher Scientific, Waltham, MA) supplemented with 25 mM HEPES, 1 mM GlutaMax, 10% FBS, and 100U/mL Penicilin-Streptomycin (Thermo Fisher Scientific, Waltham, MA). 24 hours before transfection, 2.5 × 10^6^ HeLa S3 cells were seeded in T75 plates in DMEM supplemented with 25 mM HEPES, 1 mM GlutaMax, and 10% FBS. On the day of transfection, 750 uL of Opti-MEM (ThermoFisher Scientific, Waltham, MA) was mixed with 60 uL of Lipofectamine 3000 reagent (ThermoFisher Scientific, Waltham, MA). In a separate tube was added 750 uL of Opti-MEM, 8 ug of the TDG plasmid (pCMV3-TDG), and 40 uL of the p3000 reagent, which was supplied with the Lipofectamine 3000 reagent. The two tubes were then mixed and allowed to incubate at room temperature for 15 minutes before being added to the cells. After 14 hours, cells were washed with 2 × 5 mL of PBS and treated with 3 mL of 0.25% EDTA-trypsin (ThermoFisher Scientific, Waltham, MA) at 37 °C for 5 minutes to detach cells from the plate surface. The trypsin was quenched by addition of 7 mL of DMEM and the cells were pelleted at 500 × g at 4 °C. After removing the supernatant, the cells were washed with 5 mL of PBS and pelleted at 500 x g (4 °C). Supernatant was removed and the cells were lysed in lysis buffer (85 mM KCl, 5 mM PIPES pH 8.0, 0.5% NP-40). The nuclei were collected by centrifugation at 10,000 x g (4 °C) and the resulting pellet was resuspended in RSB buffer (200 mM NaCl, 2.5 mM MgCl_2_, and 10 mM Tris-HCl pH 7.5) supplemented with 0.5% Triton X-100, 0.2% sodium deoxycholate, 0.1% SDS, and 0.05% sodium lauroyl sarcosinate. The nuclear extracts were sonicated for 6 minutes to obtain DNA fragments of approximately 500-700 bp in length, which was confirmed by native PAGE. Meanwhile, protein A Dynabeads (ThermoFisher Scientific, Waltham, MA) were pre-blocked with PBS supplemented with 0.5% BSA for 2 hours. The pre-blocked beads were then washed with 3 x 1 mL of RSB buffer supplemented with 0.5% Triton X-100 and incubated with either the IgG control or S9.6 antibody for 2 hours. For control experiments involving either the hybrid competitor (H1-C) or DNA duplex competitor (D1-C), the prepared S9.6-coated Dynabeads were further treated with 500 nM of the indicated competitor for 2 hours in RSB buffer supplemented with 0.5% Triton X-100, 0.05% sodium deoxycholate, 00.025% SDS, and 0.0125% sodium lauroyl sarcosinate. Excess competitor was removed by washing the beads with RSB buffer supplemented with 0.5% Triton X-100. The sheared nuclear extracts were then applied to the antibody-coated Dynabeads and allowed to incubate for 2 hours at 4 °C. Following immunoprecipitation, the beads were washed with 4 × 1 mL RSB buffer supplemented with 0.5% Triton X-100 and then with 2 × 1 mL RSB buffer. Bound proteins were eluted by incubating the beads with NuPAGE LDS sample buffer (ThermoFisher Scientific, Waltham, MA) supplemented with 100 mM DTT for 10 minutes at 70 °C.

### Western blotting

Eluates from the pull-down assays were resolved by 10% SDS-PAGE gels and subsequently transferred onto nitrocellulose membranes using a Trans-Blot Turbo Transfer System (Bio-Rad, Hercules, CA). Membranes were blocked with 5% non-fat milk in PSB supplemented with 0.5% tween 20 (PBST; ThermoFisher Scientific, Waltham, MA) (blocking solution) for 1 hour at room temperature on a rocker. After blocking, the membranes were washed three times with PBST and incubated overnight at 4 °C with the TDG polyclonal antibody (Proteintech, Rosemont, IL) in blocking solution. Following primary antibody incubation, the membranes were washed three times with PBST for 5 minutes and incubated with Goat anti-Rabbit IgG secondary antibody Alexa Fluor 647 (ThermoFisher Scientific, Waltham, MA) in blocking solution for 1 hour at room temperature on a rocker. The membranes were again washed three time with PBST and followed by imaging by fluorescence (Alexa Fluor 647; excitation/emission 650/671 nm) on a ChemiDoc^TM^ MP Imaging System (Bio-Rad Laboratories, Inc., Hercules, CA).

### Bioinformatics analysis

Datasets used in this study were retrieved from NCBI SRA database. They consist of TDG ChIP-Seq (GSE55660)(18), MapR and BisMapR (GSE160578)(51), H3K27ac ChIP-Seq (GSE49847; files SRR566839 and SRR566840)(52), and H3K4me3 ChIP-Seq (GSE49847; files SRR317222 and SRR317223) datasets, which were all carried out in mESCs cells.

TDG ChIP-Seq peak list was directly retrieved from GSE55660. Peak coordinates were converted from mouse genome version mm9 to mm39 (liftOver from ucsc genome browser), and blacklisted peaks were removed to end up with a final list of 71,772 peaks. Peak location was determined with the annotatePeaks.pl script from the Homer suite.(53) Fastq files retrieved from NCBI SRA database were aligned to the mm39 mouse genome version using bowtie (read length <= 50 nucleotides) or bowtie2 (read length > 50 nucleotides), and PCR duplicates were removed. Signal was retrieved at TDG ChIP-Seq (+/- 3 kb window from peak center; bin size of 10 bp) using scripts from Bedtools.(54)

Analysis of 5fc/5caC signal at TDG ChIP-Seq peaks was performed using published output files of mESC RRMAB-Seq datasets from control and shTdg mESCs (GSM1341314_RRMAB-Seq_shCtr.bed and GSM1341315_RRMAB-Seq_shTdg.bed files, from GSE55660)(18). Genomic location of CpG with reported 5fC/5caC signal was converted from mouse genome version mm9 to mm39, and 5fC/5caC signal at TDG peaks was retrieved with the intersectBed function of Bedtools.(54) Analysis was carried out using CpGs with reported 5fC/5caC signal in TDG peaks, and difference in 5fC/5caC signal between control and shTdg mESCs was determined by student t-test and considered significant if p < 0.05.

## RESULTS

### TDG binds DNA/RNA hybrid duplexes in vitro

To begin our investigation, we asked whether TDG is capable of binding to DNA/RNA hybrid duplexes in vitro. We used agarose gel-based electrophoretic mobility shift assays (EMSAs) to measure the affinity of TDG towards either DNA/DNA duplexes or DNA/RNA hybrids containing 2′-deoxy-2′-fluoroarabinouridine (U^F^), a non-cleavable substrate analogue of 2′-deoxyuridine (U) (Figure 2a). The U^F^ modification was placed within a centrally positioned CpG dinucleotide (5′-**X**pG-3′/5′-CpG-3′), which is the preferred sequence context of TDG and the most relevant biological target in context of active DNA demethylation.(45,55–57) Furthermore, two different sequences were used in order to demonstrate the generality of the observations. As shown in Figure 2b,c, TDG was indeed capable of binding to DNA/RNA hybrid duplexes H1-U^F^ and H2-U^F^ with an apparent *K*_d_ of 63 nM and 101 nM, respectively (Table 1). Surprisingly, these interactions were only ∼2–3-fold weaker than for binding to the corresponding DNA/DNA duplexes (D1-U^F^ and D2-U^F^, respectively; Figure 2b,c and Table 1), indicating that TDG can easily accommodate the non-canonical structure of the DNA/RNA hybrid. TDG formed both 1:1 and 2:1 TDG:substrate complexes with the DNA/RNA hybrids (Figure S1). Thus, in addition to binding to the U^F^ modification, a second molecule of TDG can form a non-specific complex at an adjacent unmodified site, similar to what has been reported for TDG binding to DNA/DNA duplexes.(45) Even in the absence of the U^F^ modification (H1-C and H2-C), both DNA/RNA hybrid substrates were bound tightly by TDG (Figure S2 and Table 1). This observation is consistent with TDG’s high affinity for CpG sites regardless of their modification state.(45) Taken together, these results show that TDG binds to DNA/RNA hybrid duplexes regardless of the modification state and with an affinity that is only slightly lower than for canonical DNA/DNA duplexes.

**Figure 2.**
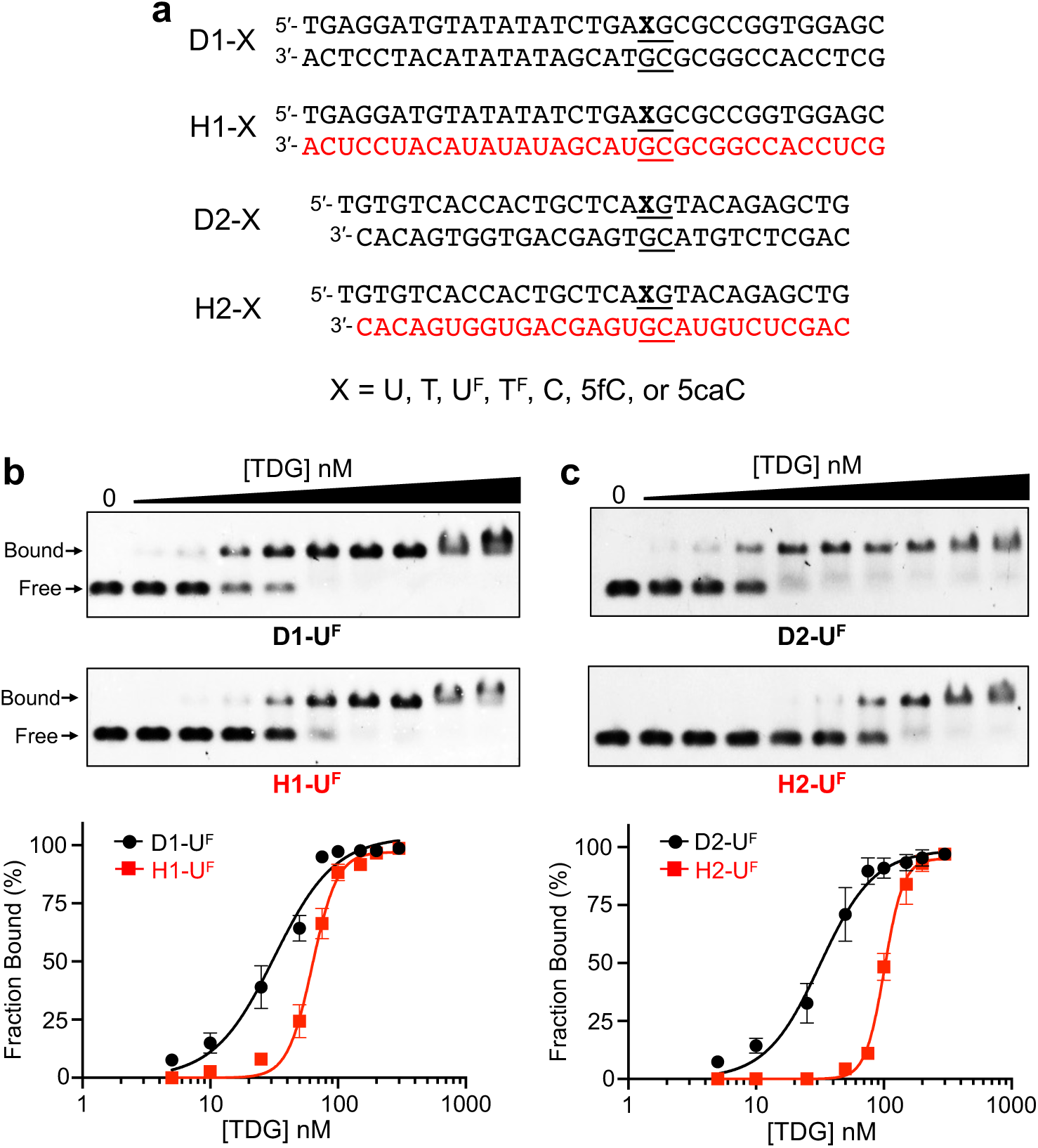
TDG binds to DNA/RNA hybrid duplexes. (a) Sequences used in this study. Black and red colors denote DNA and RNA, respectively. See also Table S1. (b,c) Representative EMSA data and corresponding saturation plots for binding of TDG to either D1/H1 (b) or D2/H2 (c). For each reaction, the indicated substrate (5 nM) was incubated with TDG (0 – 300 nM) in a buffer containing 100 mM NaCl, 2.5 mM MgCl_2_, 10 mM Tris-HCl (pH 7.5), and 5% glycerol for 30 minutes at 30 °C. Data are mean ± S.D. (n = 3).

**Table 1.** Equilibrium dissociation constants for TDG binding to the indicated substrate. 95% confidence interval (95% CI)

| Substrate | $K_d$ (nM) | 95% CI |
| --- | --- | --- |
| D1-U<sup>F</sup> | 32 | 28 – 36 |
| H1-U<sup>F</sup> | 63 | 60 – 66 |
| D2-U<sup>F</sup> | 32 | 28 – 36 |
| H2-U<sup>F</sup> | 101 | 98 – 105 |
| D1-C | 43 | 38 – 48 |
| H1-C | 67 | 60 – 74 |
| D2-C | 47 | 44 – 49 |
| H2-C | 57 | 55 – 59 |
| D1-5fC | 56 | 53 – 58 |
| H1-5fC | 60 | 57 – 64 |
| D1-5caC | 25 | 24 – 27 |
| H1-5caC | 31 | 29 – 33 |

### TDG can excise G•U pairs, but not G•T mispairs, from DNA/RNA hybrid duplexes

Having shown that TDG is capable of binding to DNA/RNA hybrids, we next asked whether it was catalytically active on these substrates. While the binding of TDG to DNA/RNA hybrids could be potentially explained by a series of nonspecific interactions, base excision requires precise positioning of the substrate within the active site of the enzyme. Therefore, we determined the rate of TDG-mediated excision of G•U or G•T pairs from DNA/RNA hybrid substrates using single-turnover kinetic experiments performed using a large excess of TDG relative to substrate (Figure S3). Single-turnover conditions are typically employed to study TDG catalysis due to strong product inhibition.(58–60) Remarkably, TDG exhibited robust excision activity for G•U pairs within both DNA/RNA hybrid duplexes tested (Figure 3a and Table 2). However, the rate of U excision from H1-U and H2-U (*k*_max_ = 1.05 min^−1^ and 0.33 min^−1^, respectively) was more than an order of magnitude slower than for the equivalent DNA/DNA duplexes (Table 2). Nevertheless, excision of U from the hybrid substrates was nearly complete within 10 minutes. In sharp contrast, TDG was nearly inactive on DNA/RNA hybrids containing a G•T mispair (H1-T and H2-T) (Figure 3b and Table 2). TDG is known to have reduced activity towards G•T mispairs compared to G•U pairs within DNA/DNA duplexes due to a steric clash between the dT methyl group and TDG that hinders flipping of the base into the enzyme active site.(61,62) Nucleotide flipping by TDG has been shown to be strongly dependent on the local environment (e.g., sequence context) and is directly related to the efficiency of base excision.(61) Thus, the inability of TDG to excise G•T mispairs from a DNA/RNA hybrid (and its reduced activity towards G•U pairs) is possibly the result of impaired nucleotide flipping when the opposing strand is RNA. This hypothesis is explored more carefully below.

**Figure 3.**
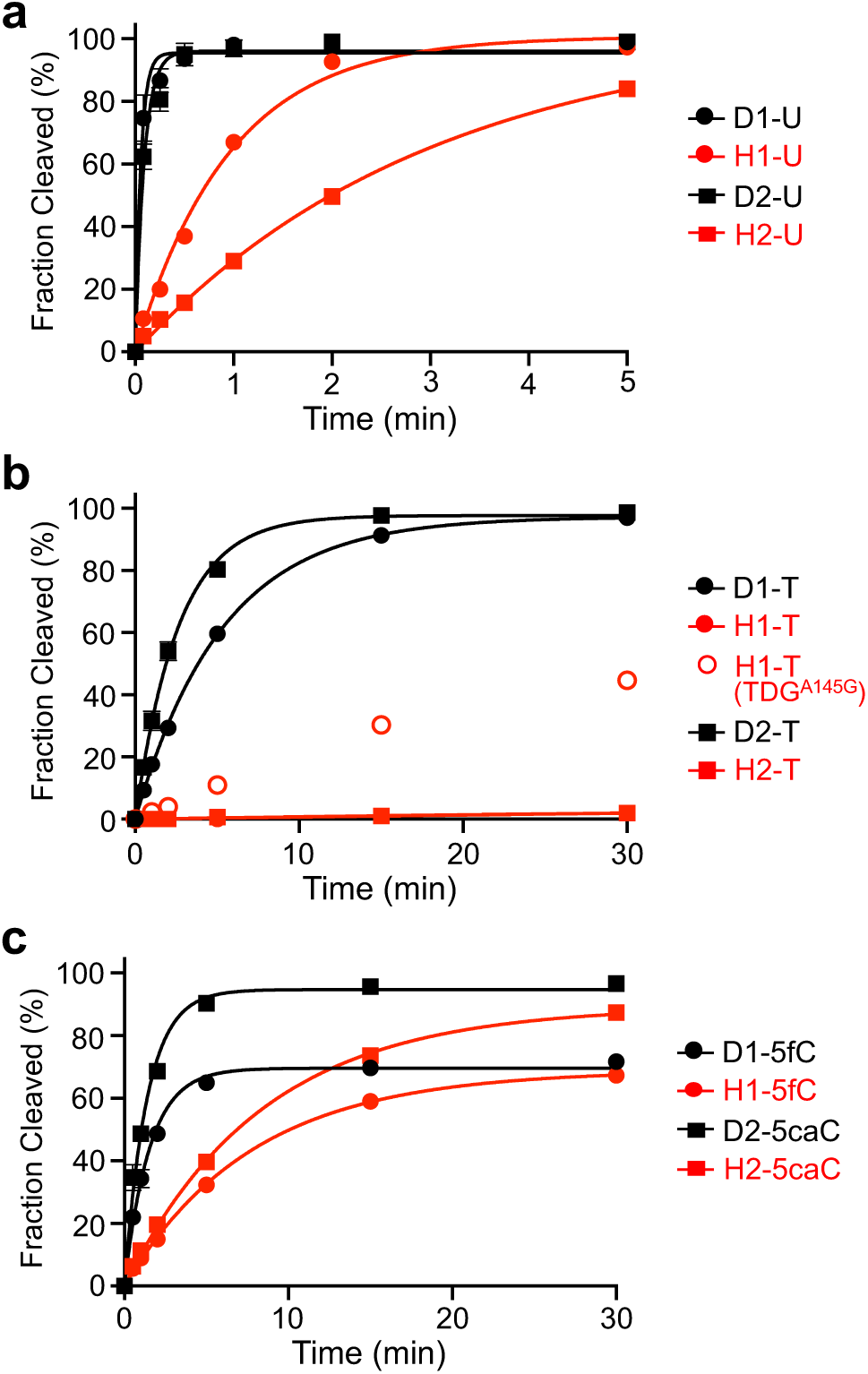
TDG is catalytically active on DNA/RNA hybrids. (a) Single-turnover kinetics of TDG (1000 nM) acting on the indicated G•U containing substrate (100 nM). (b) Single-turnover kinetics of TDG or TDG^A145G^ (1000 nM) acting on the indicated G•T containing substrate (100 nM). The data for H1-T is obscured by that for H2-T. The data for H1-T and H2-T were fitted to a linear equation. (c) Single-turnover kinetics of TDG (1000 nM) acting on the indicated 5fC/5caC containing substrate (100 nM). All reactions contained 100 mM NaCl, 2.5 mM MgCl_2_, and 10 mM Tris-HCl (pH 7.5) and were carried out at 30 °C. Data are mean ± S.D. (n = 3).

**Table 2.**
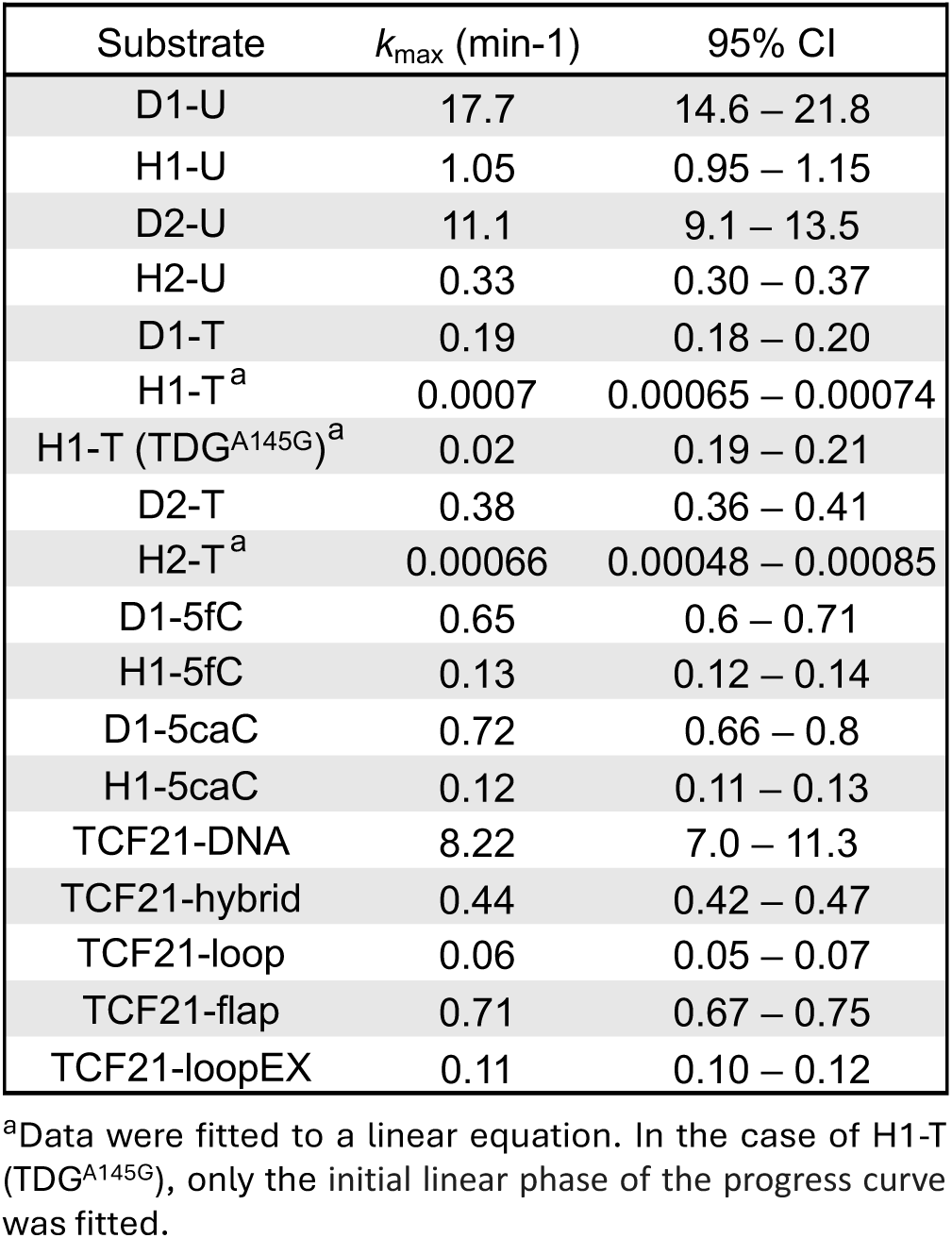
TDG excision activity (*k*_max_) for the indicated substrates at 30 °C. 95% confidence interval (95% CI)

| Substrate | $k_{\max}$ (min<sup>-1</sup>) | 95% CI |
| --- | --- | --- |
| D1-U | 17.7 | 14.6 – 21.8 |
| H1-U | 1.05 | 0.95 – 1.15 |
| D2-U | 11.1 | 9.1 – 13.5 |
| H2-U | 0.33 | 0.30 – 0.37 |
| D1-T | 0.19 | 0.18 – 0.20 |
| H1-T<sup>a</sup> | 0.0007 | 0.00065 – 0.00074 |
| H1-T (TDG<sup>A145G</sup>)<sup>a</sup> | 0.02 | 0.19 – 0.21 |
| D2-T | 0.38 | 0.36 – 0.41 |
| H2-T<sup>a</sup> | 0.00066 | 0.00048 – 0.00085 |
| D1-5fC | 0.65 | 0.6 – 0.71 |
| H1-5fC | 0.13 | 0.12 – 0.14 |
| D1-5caC | 0.72 | 0.66 – 0.8 |
| H1-5caC | 0.12 | 0.11 – 0.13 |
| TCF21-DNA | 8.22 | 7.0 – 11.3 |
| TCF21-hybrid | 0.44 | 0.42 – 0.47 |
| TCF21-loop | 0.06 | 0.05 – 0.07 |
| TCF21-flap | 0.71 | 0.67 – 0.75 |
| TCF21-loopEX | 0.11 | 0.10 – 0.12 |

<sup>a</sup>Data were fitted to a linear equation. In the case of H1-T (TDG<sup>A145G</sup>), only the initial linear phase of the progress curve was fitted.

### TDG can excise 5fC and 5caC from DNA/RNA hybrid duplexes

If R-loops are targeted for DNA demethylation in vivo, as prior work suggests (40,41), then TDG should be capable of excising oxidized cytosine derivatives from these substrates. To test this directly, we examined the capacity of TDG to excise 5fC and 5caC from CpG sites in a DNA/RNA hybrid substrate (H1-5fC and H1-5caC, respectively; Figure 2a). We note that TDG’s affinity for hybrids H1-5fC and H1-5caC was only slightly weaker than for the corresponding DNA/DNA duplexes (Figure S4 and Table 1). Using single-turnover kinetic experiments, we found that the rate of 5fC and 5caC excision from the hybrid substrates (*k*_max_ = 0.13 min^−1^ and 0.12 min^−1^, respectively) was only about 5-fold slower than from the corresponding DNA/DNA duplex (Table 2). Thus, TDG possesses substantial activity for excising 5fC and 5caC from DNA/RNA hybrids, suggesting that R-loops are compatible with TDG-mediated active DNA demethylation.

### Nucleotide flipping by TDG is impaired in DNA/RNA hybrids

The kinetic experiments above revealed that TDG has reduced glycosylase activity on DNA/RNA hybrids compared to DNA/DNA duplexes. Given that our rate constants (*k*_max_) were obtained using single turnover experiments, this observation indicated that the opposing RNA strand adversely impacted the chemical step and/or any preceding steps (e.g., conformational changes) that occur following substrate binding, which includes nucleotide flipping. Previous studies have shown that nucleotide flipping by TDG is strongly dependent on DNA context and that the flipping equilibria and glycosylase activity are positively correlated.(61) Thus, we hypothesized that the unique structural properties of the DNA/RNA hybrid may hinder nucleotide flipping, leading to the observed reduction in *k*_max_ for base excision. To examine nucleotide flipping by TDG on DNA/RNA hybrids, we employed 2′-fluoro-substituted deoxynucleotides, U^F^ and 2′-deoxy-2′-fluoroarabinothymidine (T^F^) (Figure 2a), together with ^19^F NMR. The chemical environment of the fluorine atom within U^F^ and T^F^ was previously shown to differ greatly for the stacked and flipped conformations in duplex DNA, leading to different ^19^F chemical shifts when measured by ^19^F NMR.(61) This allows the equilibrium constant for reversible nucleotide flipping (*K*_flip_) to be readily determined by comparing the integrals for peaks corresponding to the flipped (*I*_F_) and stacked (*I*_S_) states (*K*_flip_ = *I*_F_/*I*_S_).

As shown in Figure 4a, the ^19^F NMR spectrum for the DNA duplex containing a G•U^F^ pair (D2-U^F^) featured a single peak (δ^19^F −116.0 ppm), consistent with the expectation that U^F^ is predominantly stacked within the unbound duplex. This peak was lost upon the addition of TDG and a new broader peak was formed ∼6 ppm upfield (δ^19^F –122.0 ppm), which was attributed to the flipped conformation(s) of U^F^ (Figure 4b). These spectra confirmed that the vast majority of U^F^ is flipped for TDG-bound D2-U^F^ (*K*_flip_ >49) and were fully consistent with prior ^19^F NMR measurements of G•U^F^ containing DNA duplexes.(47,61) Similar to D2-U^F^, the ^19^F NMR spectrum for the DNA/RNA hybrid containing a G•U^F^ pair (H2-U^F^) showed a single peak (δ^19^F –116.8 ppm), indicating that U^F^ was also predominantly stacked within the unbound hybrid duplex (Figure 4c). This peak was shifted slightly upfield compared to the free DNA duplex D2-U^F^ (Δδ^19^F of 0.8 ppm), indicating a different chemical environment around the stacked G•U^F^ pair within the hybrid. The addition of TDG to hybrid H2-U^F^ again resulted in formation of a new upfield peak (δ^19^F –122.6 ppm) corresponding to U^F^ flipped into the TDG active site (Figure 4d). However, a small peak corresponding to the stacked G•U^F^ pair remained (δ^19^F −117.3 ppm), indicating that U^F^ was not fully flipped by TDG in the DNA/RNA hybrid. The *K*_flip_ for H2-U^F^ was calculated to be 3.0, which is at least an order of magnitude lower than for D2-U^F^ (*K*_flip_ >49). These data reveal that flipping of U^F^ from G•U^F^ pairs by TDG is impaired in DNA/RNA hybrids relative to DNA duplexes.

**Figure 4.**
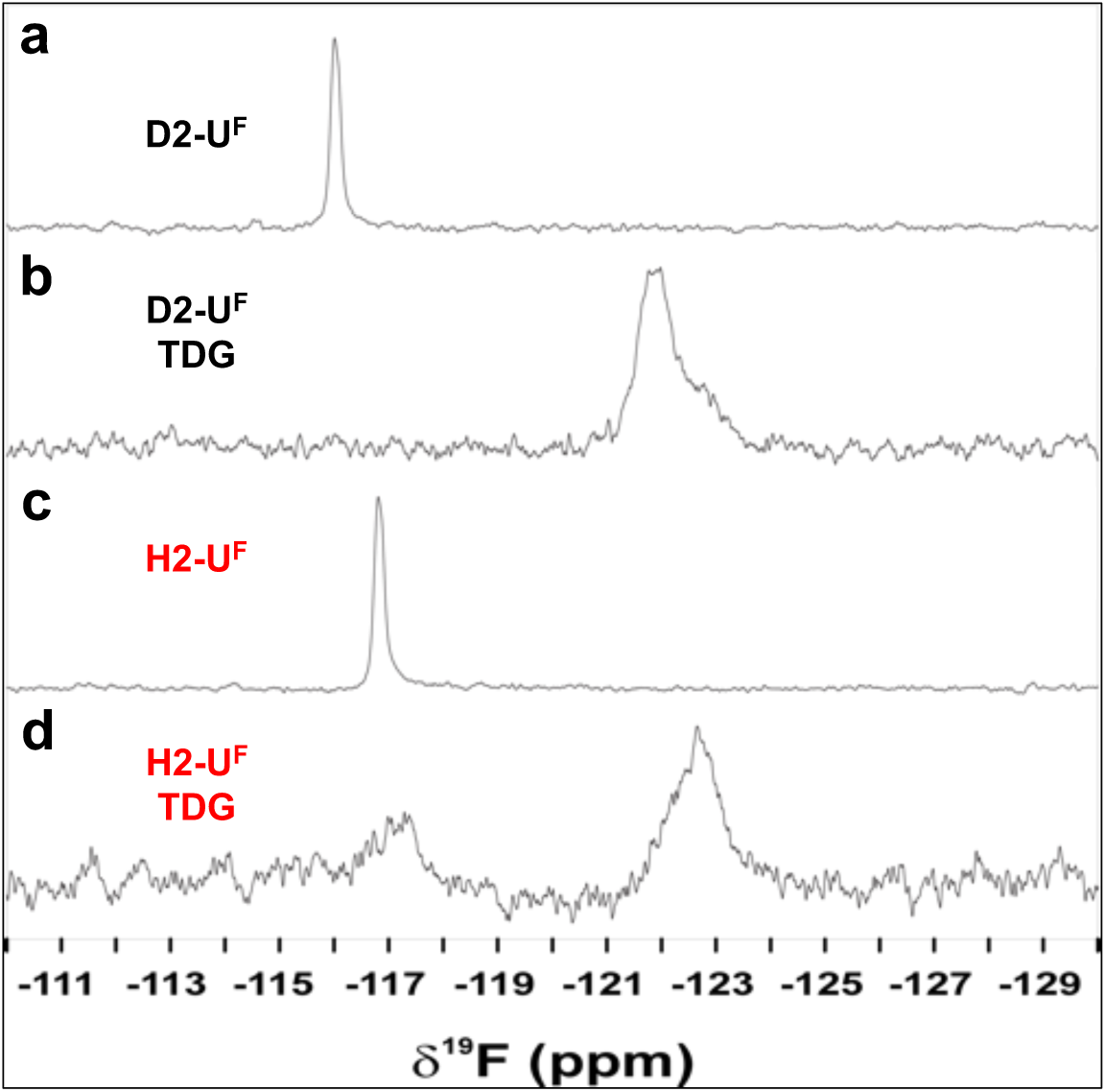
DNA/RNA hybrids impair nucleotide flipping by TDG. ^19^F NMR spectra of either DNA/DNA duplex D2-U^F^ (a,b) or DNA/RNA hybrid H2-U^F^ (c,d) in the presence or absence of TDG, collected at 25 °C. Downfield peaks (near −116 ppm) correspond to the stacked (non-flipped) conformation of U^F^, while the upfield peaks (near −122 ppm) represent the flipped conformation.

Next, we examined the excision of G•T mispairs from DNA/RNA hybrids using the A145G mutant of TDG (TDG^A145G^). Previous structural and biochemical studies have shown that Ala145 of TDG hinders nucleotide flipping of dT due to a steric clash between its methyl group and that on the thymine base.(62) Consistently, the A145G mutation greatly increases both *K*_flip_ and glycosylase activity for G•T mispairs but has little effect on G•U pairs. Thus, if the inability of TDG to excise G•T mispairs from DNA/RNA hybrids was due to hindered nucleotide flipping, we expected TDG^A145G^ to rescue this activity. Indeed, TDG^A145G^ could excise G•T mispairs from hybrid substrate H1-T, with the reaction plateauing at ∼50% after one hour (Figure 3b). Despite the relatively slow rate (*k*_obs_ = 0.02 min^−1^), this represented a nearly 30-fold increase in the rate of base excision compared to the WT enzyme on the same substrate (Table 2). Given this result, we attempted to determine *K*_flip_ for H2-T^F^ in the presence of TDG^A145G^ using ^19^F NMR as above (Figure S5). However, only a single peak (δ^19^F −117.4 ppm) corresponding to the stacked conformation of the G•T^F^ pair was observed in both the presence and absence of TDG^A145G^, suggesting that the population of flipped T^F^ is too low to be observed (< ∼2%) for H1-T^F^ bound to TDG^A145G^. While we cannot exclude the possibility that the ^19^F chemical shift for T^F^ is identical in the stacked and flipped conformations, this seems highly unlikely, given the large flipping-induced chemical shift perturbation for D2-T^F^ bound to TDG (Figure S5).

Taken together, these data strongly suggest that the reduced rate of base excision by TDG on DNA/RNA hybrids is due to impaired nucleotide flipping.

### TDG functions on authentic R-loop structures

The above experiments showed that DNA/RNA hybrids can act as substrates for TDG. However, we sought to demonstrate TDG’s ability to function on an authentic R-loop structure derived from an endogenous sequence know to be targeted for DNA demethylation. For this, we chose the promoter region of the tumor suppressor gene *TCF21*. The *TCF21* promoter was shown to form an R-loop with the lncRNA *TARID* (TCF21 antisense RNA inducing promoter demethylation), triggering local DNA demethylation and *TCF21* expression.(41) Importantly, the R-loop is positioned over several CpG sites shown to undergo demethylation in a TDG-dependent manner, suggesting that TDG can carry out base excision directly on this R-loop. We synthesized a DNA substrate consisting of nucleotides −29 to +36 (relative to the TSS) of the *TCF21* promoter and positioned an R-loop near the center (TCF21-loop) such that it covered several of the targeted CpG sites (Figure 5a and Figure S6a). To probe the potential impact of the R-loop structure on TDG activity, we also generated the corresponding “flap” (TCF21-flap) and hybrid (TCF21-hybrid) substrates (Figure 5b). TDG bound tightly to these substrates in the absence of any DNA modifications. The *K*_d_ values for the various R-loop substrates (30 – 50 nM) were similar to the DNA/DNA duplex control (TCF21-DNA, *K*_d_ = 49 nM) (Figure S6b,c). To investigate TDG’s incision activity, we incorporated a G•U pair at CpG #2 located centrally within the DNA/RNA hybrid region of these substrates and determined the rate of excision using single-turnover kinetics (Figure 5c and Table 2). Interestingly, compared to TCF21-hybrid, the rate of G•U excision from TCF21-loop was much slower (*k*_max_ is reduced 7-fold), suggesting that the presence of the ssDNA loop impeded base excision. In contrast, the rate of G•U excision from the flap substrate (TCF21-flap) was similar to the hybrid (TCF21-hybrid). TDG has been shown to bend its DNA substrate by as much as 70° upon binding (63), which is believed to help facilitate flipping of the target nucleobase into the active site for subsequent excision. Therefore, we hypothesized that the reduced rate of excision from TCF21-loop relative to TCF21-flap was due to structural constraints imposed by the DNA loop that impeded TDG’s ability to bend the substrate. Consistently, increasing the size of the ssDNA loop from 15-nt to 30-nt (TCF21-loopEX) led to a ∼2-fold increase in base excision rate (Figure 5c and Table 2). Together, these results demonstrate that TDG is able to bind and excise authentic R-loop structures and suggest that TDG’s activity on R-loops could be dependent on the length and structure of the ssDNA loop.

**Figure 5.**
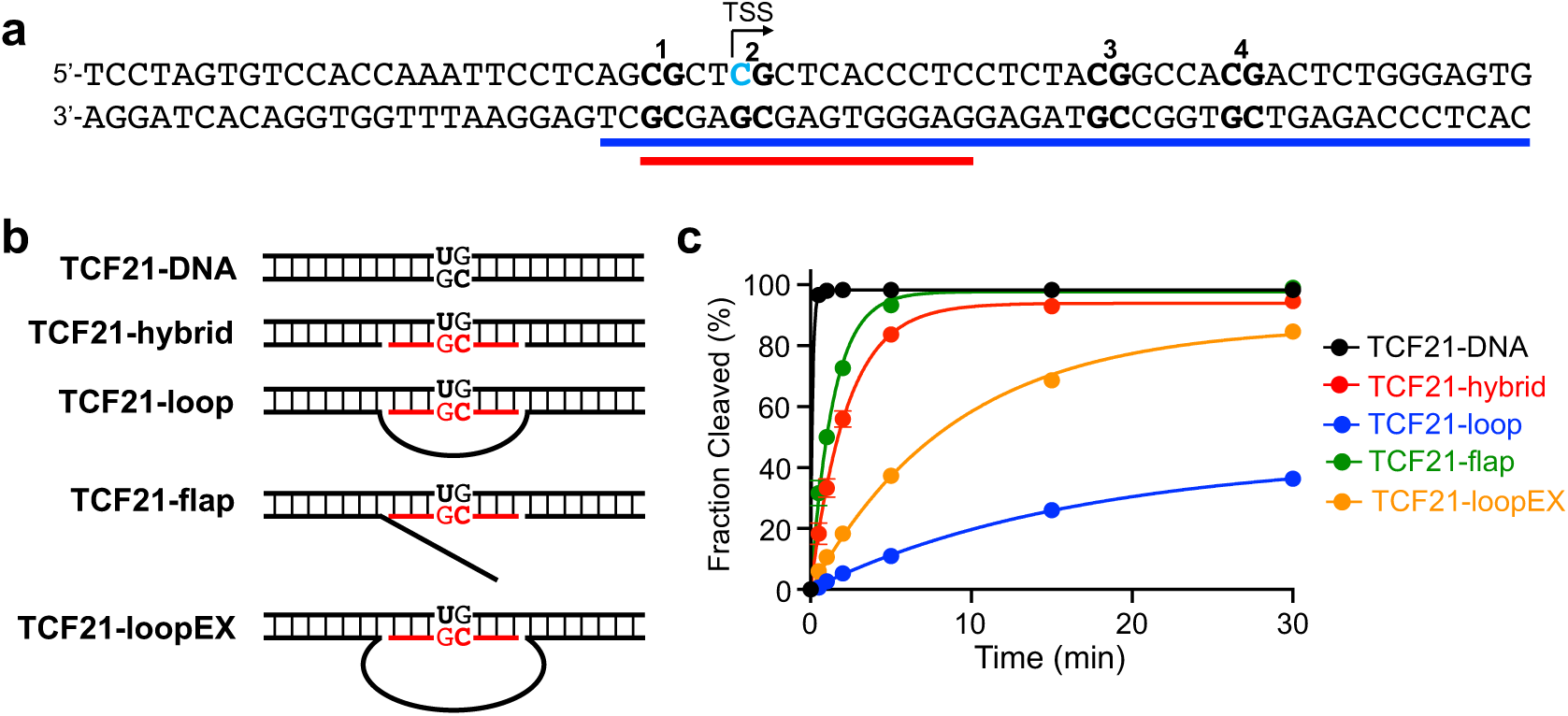
TDG functions on authentic R-loop structures. (a) DNA used in this work. The sequence is derived from the *TCF21* gene promoter (−29 to +36 relative to the TSS). The blue line indicates the position of the endogenous R-loop, whereas the red line indicates the position of the R-loop used herein. Individual CpG dinucleotides are numbered and the position of dU incorporation is indicated by blue text. See Figure S6 for sequence details. (b) Schematic illustration of the *TCF21*-derived substrates. Black and red colors denote DNA and RNA, respectively. (c) Single-turnover kinetics of TDG (1000 nM) acting on the indicated substrate (100 nM). Reaction conditions are identical to those described in Figure 3. Data are mean ± S.D. (n = 3).

### R-loops direct the strand selectivity of TDG and prevent DSB formation at symmetrically modified CpG

Active DNA demethylation has been shown to occur in a strand-selective fashion at transcriptionally active promoters. (22,64) Furthermore, genome-wide mapping studies of 5fC and 5caC suggest that TDG’s processivity for these modifications is strand selective at some CpGs. However, it remains unclear how this selectivity is achieved, especially considering that TDG has no apparent strand preference when presented with symmetrically modified CpGs in vitro.(7) In light of our findings above, we reasoned that the strand selectivity of TDG could be influenced by R-loops. Because TDG is catalytically inactive on ssDNA(65), only the DNA strand that is hybridized to the RNA is expected to be a substrate for base excision. Thus, for CpGs within an R-loop, TDG’s activity should be directed towards the DNA strand contained within the DNA/RNA hybrid, even if both sides of the CpG contain a potential substrate. To test this directly, we assembled a 60-bp DNA substrate in which the central CpG site was symmetrically modified with G•U pairs (symGU) (Figure 6a and Table S1). The top and bottom strands of symGU were labelled with Cy5 and Cy3, respectively, allowing for base excision on both strands to be monitored simultaneously within the same reaction. As shown in Figures 6a and S7, TDG acted evenly on both strands of symGU under single-turnover conditions. The amount of single-strand incision rapidly reached 50% for both strands before starting to plateau. This observations is consistent with prior results showing that the processing of one side of a CpG largely inhibits processing of the other, likely due to the tight interaction of TDG with the AP-site product.(7) Nevertheless, the extent of excision reached >80% for both strands after 30 minutes. In stark contrast, formation of a 15-nt long R-loop over the symmetrically modified CpG (symGU-R) resulted in almost exclusive base excision from the top strand (i.e., the DNA/RNA hybrid) (Figure 6b and Figure S7). The rate of excision from the top strand (*k*_max_ = 2.0 min^−1^) was >700-fold faster than from the bottom, unhybridized strand (*k*_max_ = 0.0027 min^−1^). These results demonstrate that R-loops can effectively direct the strand selectivity of TDG.

**Figure 6.**
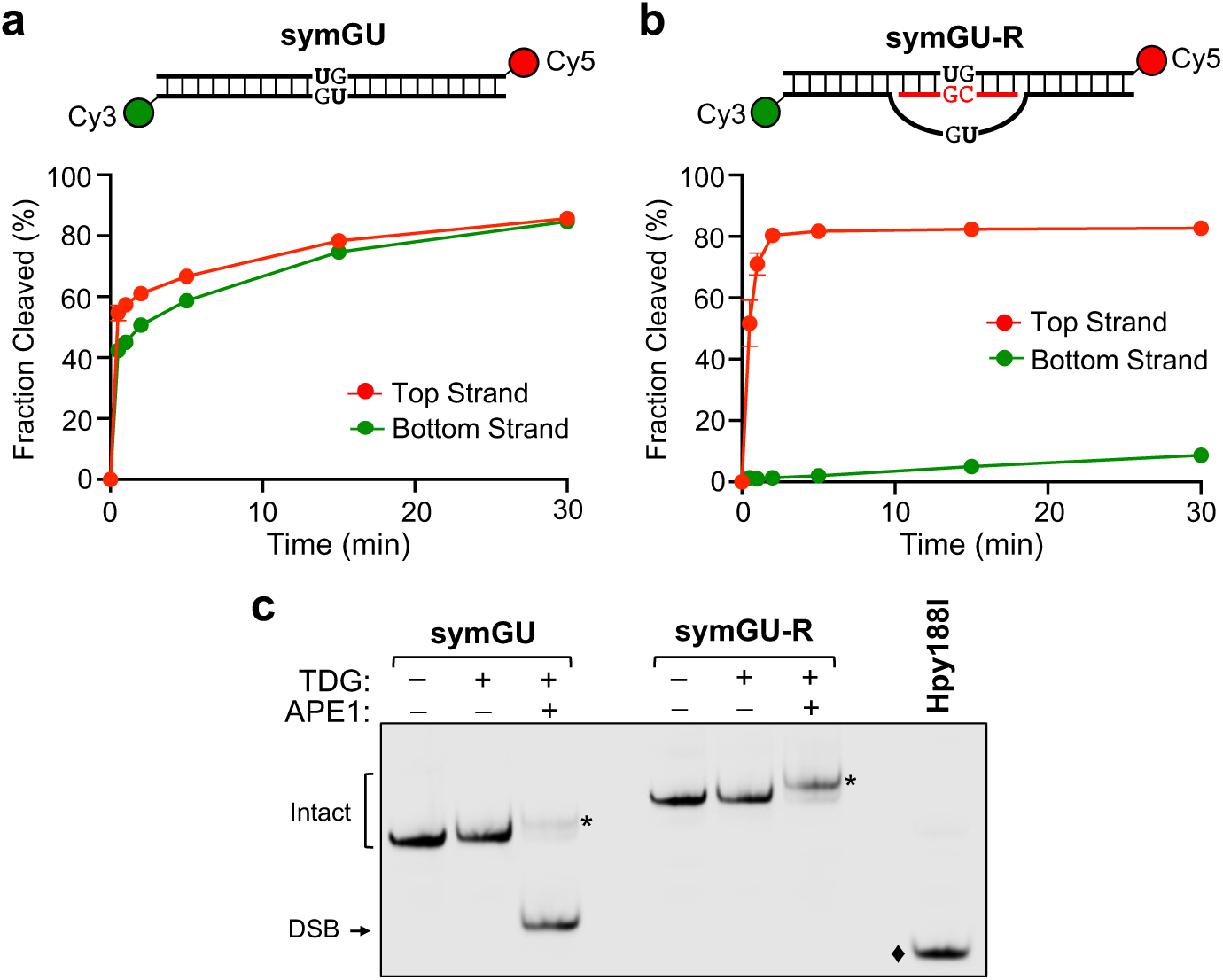
Processing of symmetrically modified R-loops by TDG. (a,b) Single-turnover kinetics of TDG (1000 nM) acting on either (a) symGU or (b) symGU-R (100 nM). Substrates were labelled on the top strand with Cy5 (red) and on the bottom strand with Cy3 (green). Reaction conditions are identical to those described in Figure 3. Data are mean ± S.D. (n = 3). (c) Representative native PAGE gel showing the formation of DSBs. The indicated substrate (100 nM) was treated with TDG (200 nM) and/or APE1 (20 nM) in a buffer containing 100 mM NaCl, 2.5 mM MgCl_2_, and 10 mM Tris-HCl (pH 7.5) for 30 minutes at 30 °C. Asterisks indicated the nicked duplex. The diamond indicates a DSB control product generated via the treatment of symGU with restriction enzyme Hpy188I.

Base excision of symGU exceeded 50% on both strands under the conditions used herein (Figure 6a), indicating that TDG-initiated BER could induce the formation of DNA double-strand breaks (DSBs) at CpGs that are symmetrically modified with its nucleobase substrates. Indeed, when monitored by native gel electrophoresis (Figure 6c), the combined action of TDG and APE1 produced a substantial fraction of DSBs from DNA symGU (>80%). Similar observations have been observed at CpGs symmetrically modified with 5caC.(7) However, no detectable DSBs were produced upon incubation of the R-loop substrate symGU-R with TDG and APE1 (Figure 6c), consistent with the almost exclusive processing of the top, hybrid strand by the two enzymes. We note that efficient incision of DNA AP sites by APE1 in DNA/RNA hybrids has been previously reported.(66) Thus, in addition to directing the strand selectivity of TDG at symmetrically modified CpG substrates, the formation of R-loops may help avoid the formation of cytotoxic DSBs during active DNA demethylation.

### TDG interacts with R-loops in cells

Genome-wide mapping studies of TDG occupancy show that TDG is enriched at promoters and enhancers of active genes, a distribution similar to R-loops.(18,33,51,67,68) To investigate this potential overlap more carefully, we compared the genome-wide distribution of TDG binding sites and R-loops in mouse embryonic stem cells (mESCs) using published TDG chromatin immunoprecipitation sequencing (ChIP-seq)(18) and MapR(51) datasets, respectively. MapR employs a catalytically inactive RNase H to target micrococcal nuclease to R-loops to cleave and release them for high-throughput sequencing. As shown in Figure 7a and S8, TDG binding sites were strongly enriched for MapR signals. Of the 71,772 TDG ChIP-seq peaks analysed, 21,602 of them (30.1%) overlapped with a MapR peak (i.e., an R-loop). Overlap was strongest at gene promoters (referred to as TDG/R-loop promoters; 48.4%, TSS −1 kb to +100 bp), with overlapping peaks in these regions being highly enriched for marks of active transcription (H3K27ac and H3K4me3) (Figure 7a-c). Given that 5fC and 5caC are also enriched at the promoters of actively transcribed genes(18,19), we asked whether TDG/R-loop promoters undergo active DNA demethylation by comparing overlapping TDG ChIP-seq and MapR peaks with genome-wide maps of 5fC and 5caC generated using control and TDG-silenced mESCs.(18) Indeed, we found that 5fC and 5caC levels at TDG/R-loop promoters increased upon depletion of TDG (Figure 7d), consistent with an active TDG-dependent 5fC and 5caC excision mechanism taking place at these sites. Taken together, these data demonstrate that TDG localizes to sites of R-loop formation in vivo, where it is capable of excising 5fC and 5caC.

**Figure 7.**
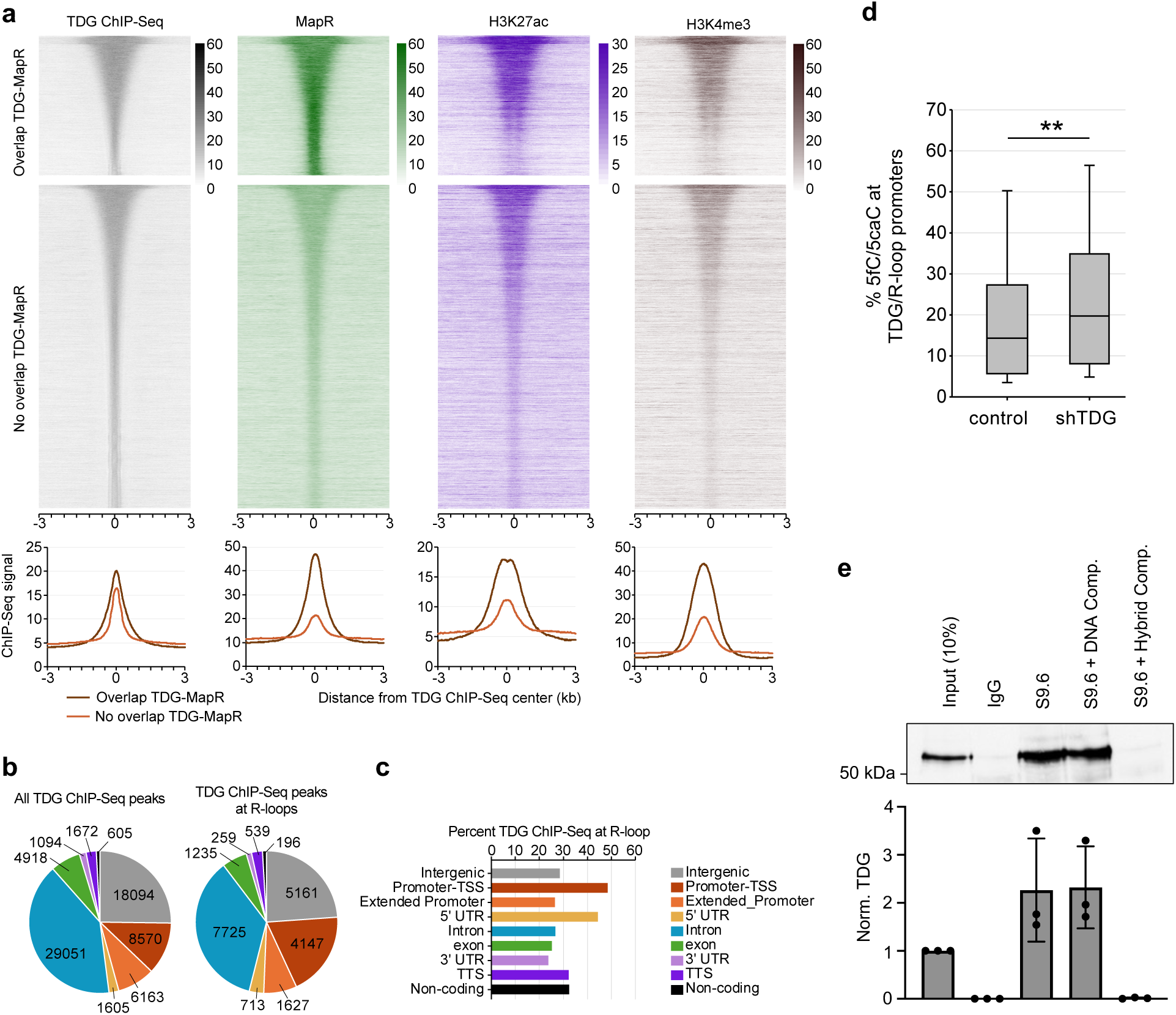
TDG interacts with R-loops in mESCs. (a) Heatmap representations on a window of ±3 kb around the center of TDG ChIP-seq peaks. Reads were parsed based on the overlap of TDG peaks with MapR peaks and ordered based on TDG ChIP-Seq signal. (b) Pie charts comparing the genome-wide distribution of all TDG peaks (n = 71,772) to those that overlap with R-loops (n = 21,602). Promoter-TSS: +/-1 kb from TSS; Extended promoter: −10 kb to −1 kb from TSS. (c) Percent of TDG peaks at R-loops. (d) Box plot of the percentage of total 5fC/5caC at TDG/R-loop promoters in TDG knockdown (shTDG) versus control mESCs. **p < 0.01. (e) TDG is associated with cellular R-loops. Lysates from TDG-expressing HeLa cells were immunoprecipitated with the S9.6 antibody and co-precipitated TDG was visualized by Western blot. Where indicated, the S9.6 antibody was treated with competitor (comp.) prior to the immunoprecipitation step. Quantification of precipitated TDG from three independent DRIP experiments is shown below. Data are mean ± S.D. normalized to the input (10%).

To further confirm that TDG interacts with R-loops in cells, we carried out DNA-RNA immunoprecipitation (DRIP) using the R-loop-specific antibody S9.6 and monitored the co-precipitation of TDG by Western blotting.(40) The assay was carried out using HeLa cells that were transfected with a plasmid encoding full-length human TDG. As shown in Figure 7e, TDG was strongly enriched following immunoprecipitation (IP) of the cell lysates with the S9.6 antibody compared to the IgG control. Furthermore, incubation of the S9.6 antibody with a synthetic R-loop competitor (but not DNA duplex competitor) prior to the IP step abolished its ability to precipitate TDG, confirming that the assay specifically enriched R-loop bound TDG. The results of the DRIP assay, together with our observation that a large portion of TDG ChIP-seq peaks overlap with genomic R-loops (Figure 7a), strongly support the interaction of TDG with R-loops in cells.

## DISCUSSION

Originally thought to be rare byproducts of transcription, R-loops have now been shown to play a key role in a variety of nuclear processes. In particular, the number of connections between R-loop formation and chromatin modifications continues to grow. Studies now indicate that R-loops may act as an epigenetic mark, being read by chromatin remodels and other proteins to effect changes in chromatin state.(69) Notably, R-loops have been shown to promote local DNA demethylation by recruiting associated proteins, including TETs and TDG.(41) Herein, we showed for the first time that TDG, a central component of the active DNA demethylation machinery, binds tightly to R-loops *in vitro* and can excise oxidized cytosines from DNA/RNA hybrid duplexes. Furthermore, we demonstrated that R-loops can direct the strand selectivity of TDG at CpG dinucleotides, providing a potential explanation for the strand-specific distribution of 5fC/5caC observed at gene promoters.(22,64) Finally, our data suggest that interactions between R-loops and TDG may be extensive in human cells. Together, our results provide critical support for the role of R-loops in regulating DNA demethylation and offer a mechanism whereby 5fC/5caC are directly excised from DNA/RNA hybrids.

DNA duplexes adopt a B-form helix whereas DNA/RNA hybrids favor A-form.(70–74) Despite these structural differences, our data show TDG can accommodate both types of substrates, but how? A potential explanation lies in the intrinsic structural plasticity of the DNA/RNA hybrid. While DNA/RNA hybrids have many characteristics of the A-form, NMR studies and molecular dynamic simulations indicate that they exist as an ensemble of conformation having both B-like and A-like structures.(70–74) TDG may exploit this structural plasticity to sculpt the hybrid to fit its DNA-binding pocket. Indeed, the ability of TDG and other DNA glycosylases to conformationally distort their substrates (e.g., through DNA bending and pinching) via protein-DNA interactions is well document.(56,63,75,76) The catalytic pocket of TDG has also been shown to be adaptable, which could further facilitate its interactions with the hybrid. For example, cis–trans isomerization of proline 155 allows for large conformations changes to occur near the active site.(44) Regardless of how TDG interacts with DNA/RNA hybrids, our ^19^F NMR studies reveal that the resulting complex differs from that made between TDG and duplex DNA and leads to impaired nucleotide flipping. Flipping of the target nucleobase into the TDG active site and stabilization of its extrahelical conformation requires a precise network of interactions with the substrate, which may simply be more difficult to achieve with the hybrid (e.g., due to steric clash). The unique thermodynamic and mechanical properties of the hybrid relative to duplex DNA are also expected to contribute to flipping. The thermal stability of DNA/RNA hybrids is generally greater than for duplex DNA, making the hybrid less prone to base flipping.(77–79) Consistently, the ϕλG°_37_ values for the unmodified hybrid sequences H1-C (ϕλG°_37_ = −44.50 kcal mol^−1^) and H2-C (ϕλG°_37_ = −39.50 kcal mol^−1^) used herein were calculated to be 2.2 kcal mol^−1^ and 2.45 kcal mol^−1^ less than their corresponding DNA duplexes, respectively, using reported nearest neighbour parameters (in 1 M NaCl).(79,80) Furthermore, theoretical studies predict that DNA/RNA hybrids are stiffer and more resistant to local deformation than DNA duplexes, potentially making it more difficult to be bent by TDG.(70) Bending of the substrate is believed to facilitate flipping of the target nucleobase into TDG’s active site for subsequent excision.(63) Ultimately, addressing questions about how TDG binds to and excises nucleobases from DNA/RNA hybrids will require future structural studies.

Because R-loops are formed in a sequence-specific manner, they provide an attractive mechanism for the precise targeting of CpGs for active DNA demethylation. Indeed, there is increasing evidence that R-loops act as guides to recruit proteins involved in DNA demethylation, including GADD45A, TET1, and TDG, to specific gene promoters.(25,41,69) By showing that TDG binds to R-loops and excises 5fC and 5caC from DNA/RNA hybrids, our study provides critical support for this R-loop targeting model. However, our results suggest additional roles for R-loops during DNA demethylation that go beyond recruiting. For instance, R-loops are inhibitory for nucleosome formation and are typically associated with increased chromatin accessibility.(81) We previously showed that nucleosomes and folded chromatin fibers impede DNA binding and base excision by TDG.(60) Thus, R-loop formation may be important for providing TDG access to the underlying DNA. Additionally, our results demonstrate that R-loops can target TDG-mediated base excision to the DNA strand contained within the DNA/RNA hybrid, even if both sides of the CpG contain a potential substrate. This provides a straightforward solution for how TDG achieves strand-selective processing of 5fC/5caC at symmetrically modified CpG. This result also raises the intriguing possibility that R-loops can direct the strand selectivity of TETs, which would make R-loops a key determinant for choosing which strand undergoes active DNA demethylation at symmetrically methylated CpGs. Although TET proteins have been shown to oxidize 5mC within DNA/RNA hybrids (82), whether R-loops influence their strand selectivity remains unknown and warrants further investigation. Finally, our data suggest that R-loops play a protective role during active DNA demethylation by preventing the formation of DSBs. Most CpG dinucleotides in mammals are symmetrically methylated, generating a high potential for DNA DSBs upon demethylation and subsequent BER. Previous studies proposed that DSB are mostly avoided due, in part, to the high affinity of TDG for its AP-site product, which prevents the opposite strand from being processed simultaneously.(7) However, APE1 has been shown to stimulate the catalytic turnover of TDG by disrupting the product complex,(83) suggesting that tight coupling of TDG and APE1 with the downstream BER machinery is necessary to avoid the formation of DSBs. This is highlighted by our observation that the combined activities of TDG and APE1 yielded significant DSBs in the absence of downstream BER enzymes (Figure 6c). In contrast, no DNA DSBs were detected with R-loop substrates, which is attributed to the weak activity of TDG and APE1 on the looped out ssDNA.(84,85) Thus, by essentially masking one side of the CpG from the BER machinery, the formation of R-loops provides a simple mechanism for avoiding DSBs during active DNA demethylation. It is worth noting here that TDG was found to be weakly active on G•T mispairs in DNA/RNA hybrids. This indicates that G•T mispairs, which arise frequently at CpG dinucleotides due to the higher rate of deamination of 5mC compared with unmethylated cytosine (86), will avoid repair as the result of R-loop formation. In this scenario, the formation of R-loops could be considered a disadvantage by promoting C to T mutations within CpG dinucleotides.

In conclusion, our data provide proof of TDG functionality on R-loops and suggest several novel regulatory roles for R-loops (and RNA in general) during active DNA demethylation. This new paradigm for TDG has important implications for all TDG-dependent processes and provides several exciting avenues for future investigation.

## DATA AVAILABILITY

The data generated during all experiments is available from the author upon reasonable request.

## SUPPORTING INFORMATION

This article contains supporting information. Figures S1 – S19 and Table S1.

## ACKNOWLEDGEMENTS

The authors acknowledge the Structural Biology Shared Service of the University of Maryland Marlene and Stewart Greenebaum Comprehensive Cancer Center (UMGCCC) for NMR spectroscopy. Equipment used for this work was supported in part by the Maryland Department of Health Cigarette Restitution Fund Program (CH-649-CRF) and a National Cancer Institute Cancer Center Support Grant (P30CA134274). Portions of this research were conducted with the advanced computing resources provided by Texas A&M High Performance Research Computing.

