## Supplemental Information for "Thymine DNA Glycosylase Binds to R-Loops and Excises 5-Formyl and 5-Carboxyl Cytosine from DNA/RNA Hybrids"

### S1. Supplementary Figures

#### Figure S1

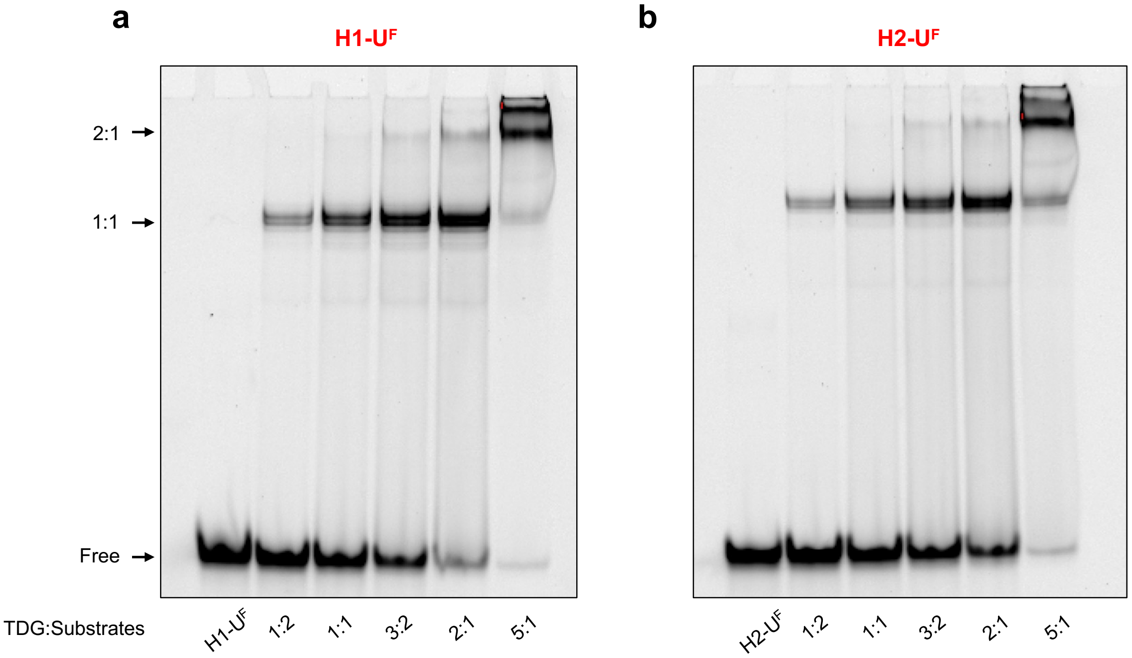


**Figure S1.** Representative native PAGE gels (10%, 29:1 acrylamid:bisacrylamide) showing the formation of 1:1 and 2:1 complexes of TDG (100 – 1000 nM) with either (a) H1-$U^F$ or (b) H2-$U^F$. For each reaction, the indicated substrate (200 nM) was mixed with TDG in a buffer containing 100 mM NaCl, 2.5 mM $MgCl_2$, and 10 mM Tris-HCl (pH 7.5), and was incubated at 30 °C for 30 minutes. Uncropped gel images are presented in Figure S15.

**Figure S2**

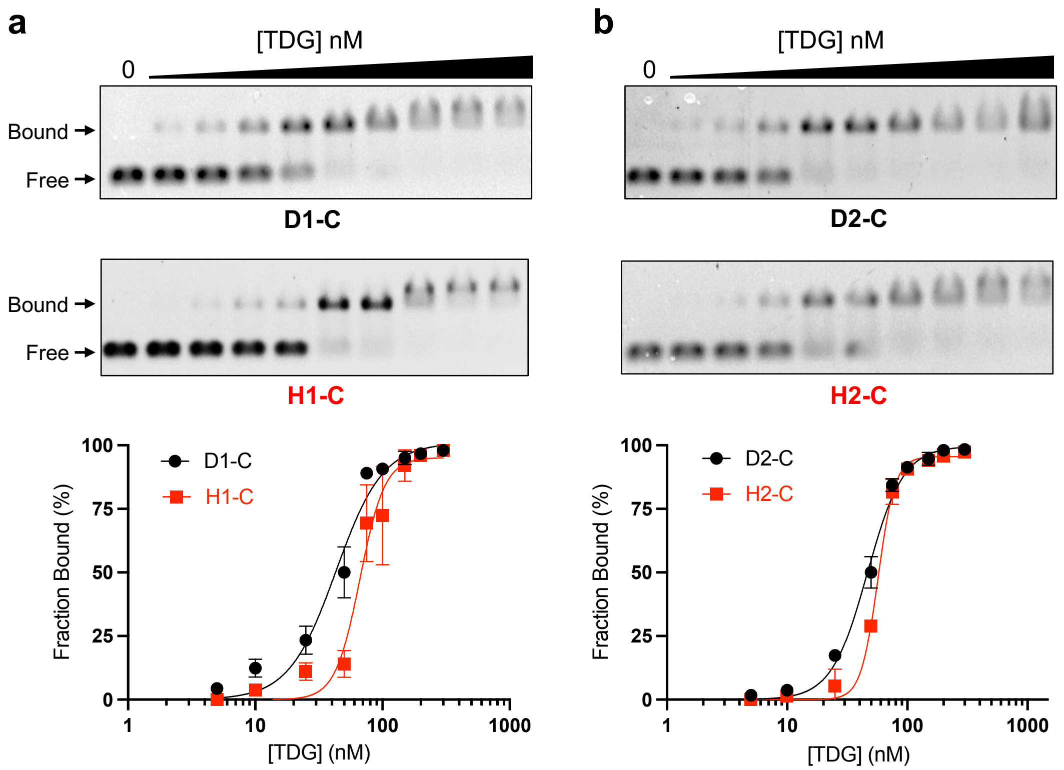


**Figure S2.** (a) Representative EMSA data and corresponding saturation plots for binding of TDG (0 – 300 nM) to either D1-C or H1-C (5 nM). (b) Representative EMSA data and corresponding saturation plots for binding of TDG (0 – 300 nM) to either D2-C or H2-C (5 nM). Reaction conditions are the same as those described in Figure 2b,c. Data are mean ± S.D. (n = 3). Uncropped gel images are presented in Figure S16.

**Figure S3**

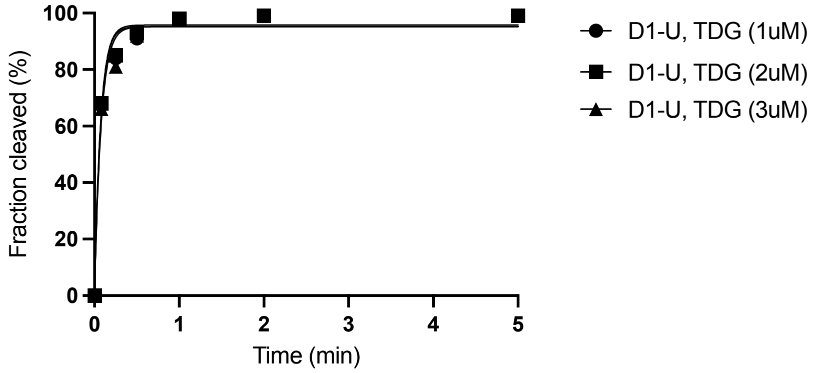


**Figure S3.** Single-turnover kinetics using different concentrations of TDG (1000, 2000, and 3000 nM) acting on D1-U (100 nM). For each reaction, D1-U (100 nM) was mixed with TDG (1000, 2000, and 3000 nM) respectively in a buffer containing 100 mM NaCl, 10 mM Tris-HCl pH 7.5, 2.5 mM $MgCl_2$ and was incubated at 30 °C for 30 minutes. Data are mean ± S.D. (n = 3).

**Figure S4**

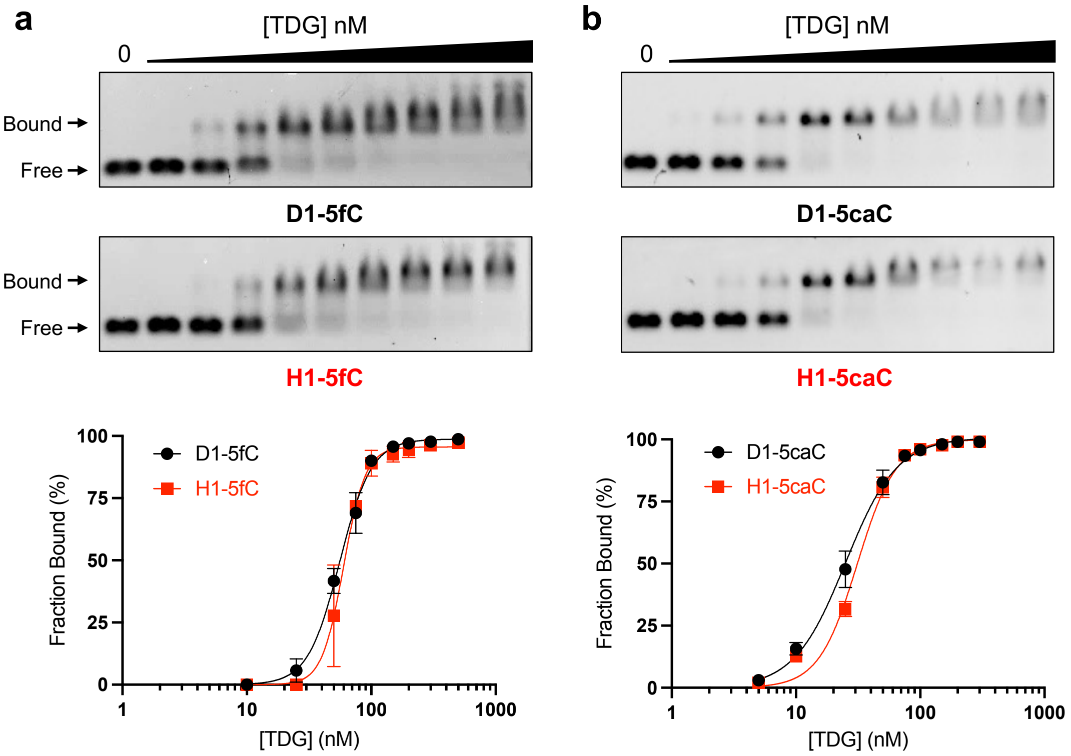


**Figure S4.** (a) Representative EMSA data and corresponding saturation plots for binding of TDG (0 – 300 nM) to either D1-5fC or H1-5fC (5 nM). (b) Representative EMSA data and corresponding saturation plots for binding of TDG (0 – 300 nM) to either D2-5caC or H2-5caC (5 nM). Reaction conditions are the same as those described in Figure 2b,c. Data are mean ± S.D. (n = 3). Uncropped gel images are presented in Figure S17.

**Figure S5**

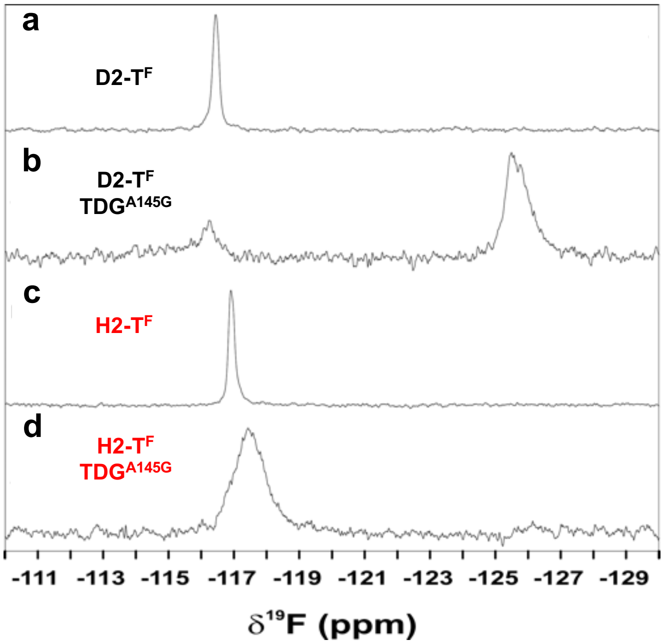


**Figure S5.** $^{19}$F NMR spectra for D2-$T^F$ (a,b) or H2-$T^F$ (c,d) in the absence or presence of $TDG^{A145G}$. The DNA concentrations were 67 μM to 75 μM and the enzyme concentration was at least twofold greater than DNA.

### Figure S6

a

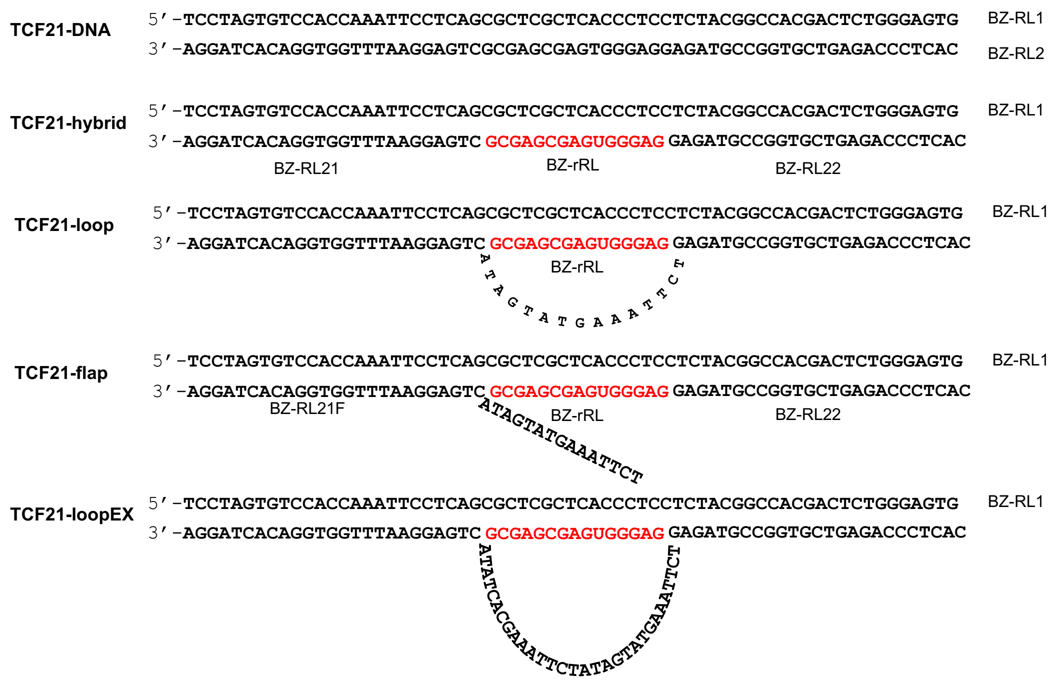


b

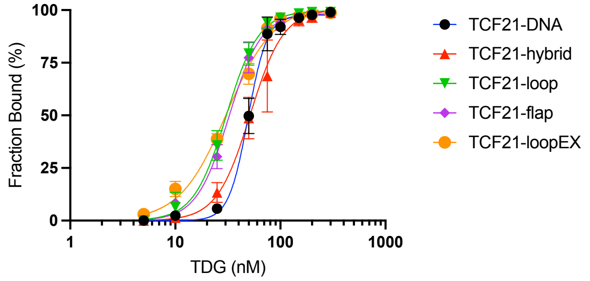


c

| Substrate | $K_d$ (nM) | 95% CI |
|---|---|---|
| TCF21-DNA | 49 | 47 - 51 |
| TCF21-hybrid | 52 | 47 - 52 |
| TCF21-loop | 31 | 29 - 33 |
| TCF21-flap | 33 | 31 - 35 |
| TCF21-loopEX | 31 | 29 - 33 |

**Figure S6.** (a) Schematic and sequences of all TCF21-derived substrates. Black and red colors denote DNA and RNA, respectively. (b) Saturation plots for binding of TDG (0 – 300 nM) with the indicated R-loop substrates. Reaction conditions are the same as those described in Figure 2b,c. (c) Equilibrium dissociation constats for TDG binding to the indicated substrates. 95% confidence interval (95% CI).

**Figure S7**

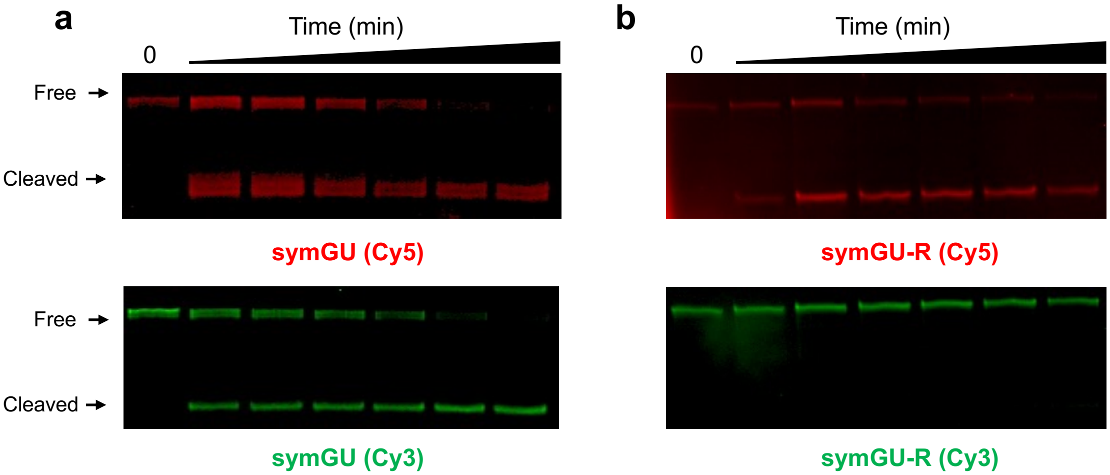


**Figure S7.** Single-turnover kinetics of TDG-mediated cleavage of symGU (a) or symGU-R (b) as measured by denaturing PAGE (20%, 19:1 acrylamide:bisacrylamide). Representative gels are shown, which were scanned using the indicated channel. Reaction conditions are the same as those described in Figure 3. Uncropped gel images are presented in Figure S18.

**Figure S8**

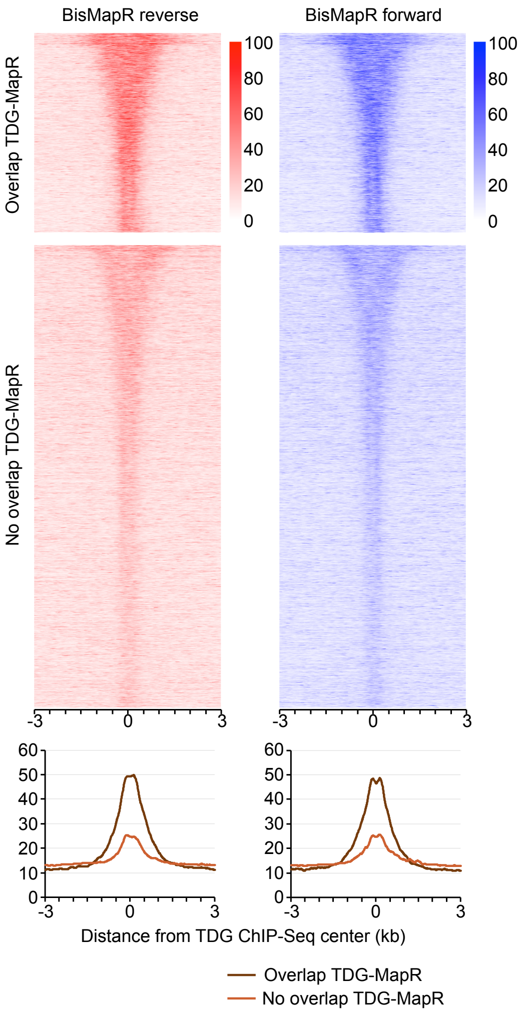


**Figure S8.** TDG ChIP-Seq signal is enriched at R-loops. (top) Heatmap representations of BisMapR reverse and forward signal on a window of ±3 kb around the center of TDG ChIP-seq peaks. Reads were parsed based on the overlap of TDG peaks with MapR peaks and ordered based on TDG ChIP-Seq signal, as in Figure 7a. (bottom) Quantification of BisMapR reverse and forward signal signal at TDG peaks that overlap with an R-loop (brown) or not (orange).

**Figure S9**

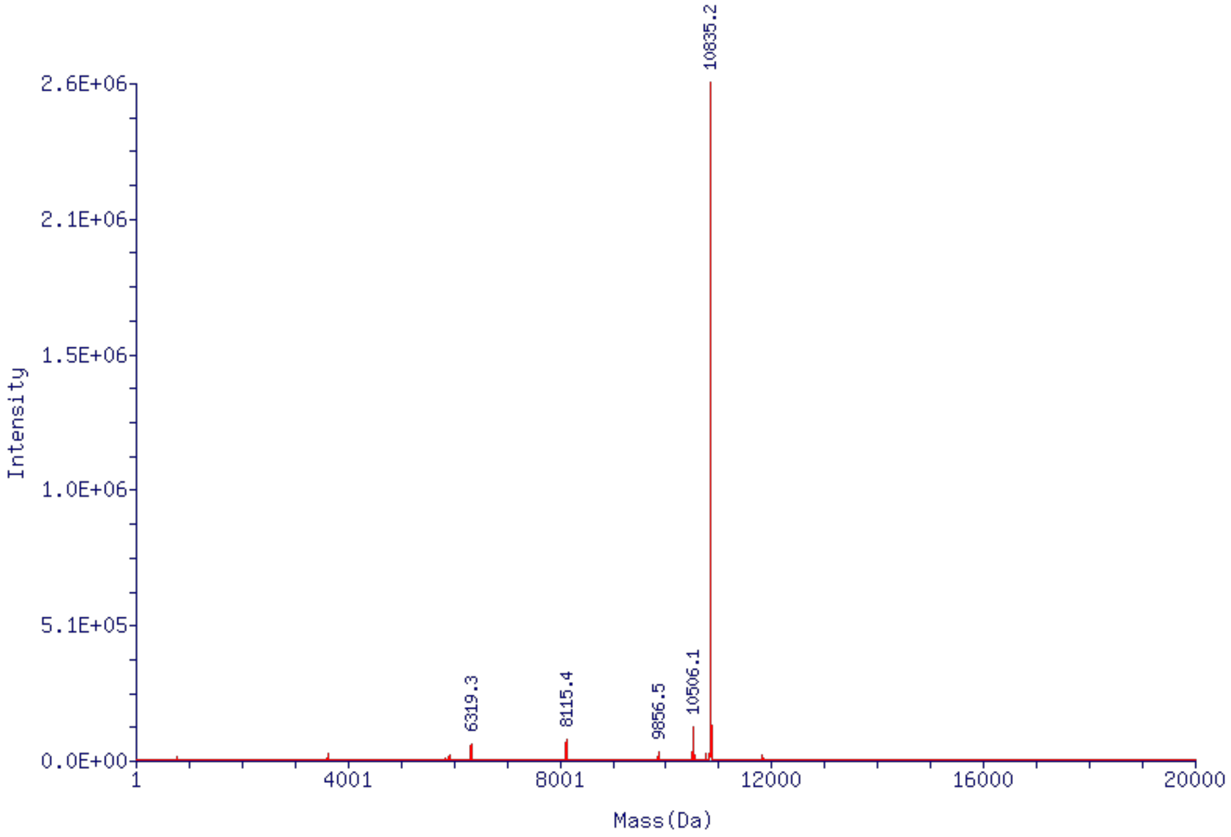


**Figure S9**. ESI-MS of BZ-dU$^{F}$FWD. Mass calculated: 10835.14 Da; Mass found: 10835.2 Da.

**Figure S10**

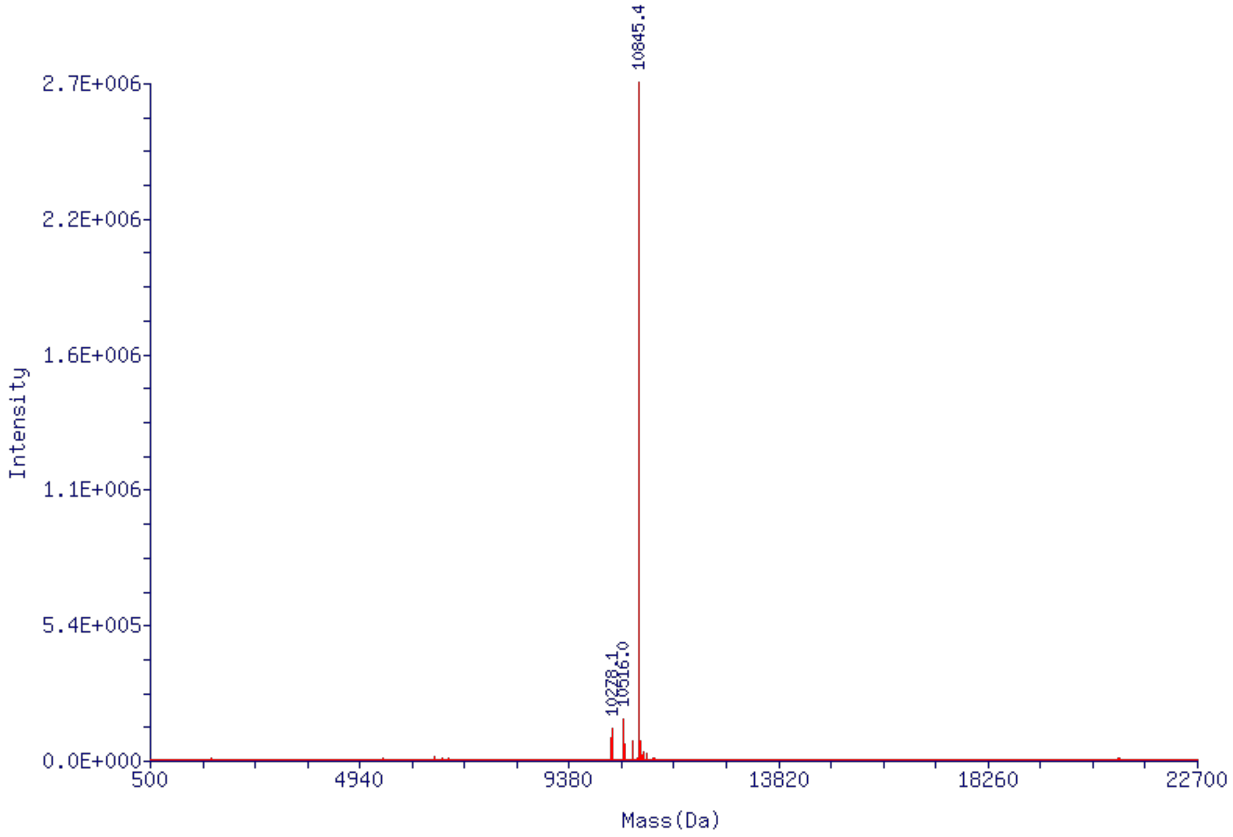


**Figure S10**. ESI-MS of BZ-5fCFWD. Mass calculated: 10844.17 Da, Mass found: 10845.4 Da.

**Figure S11**

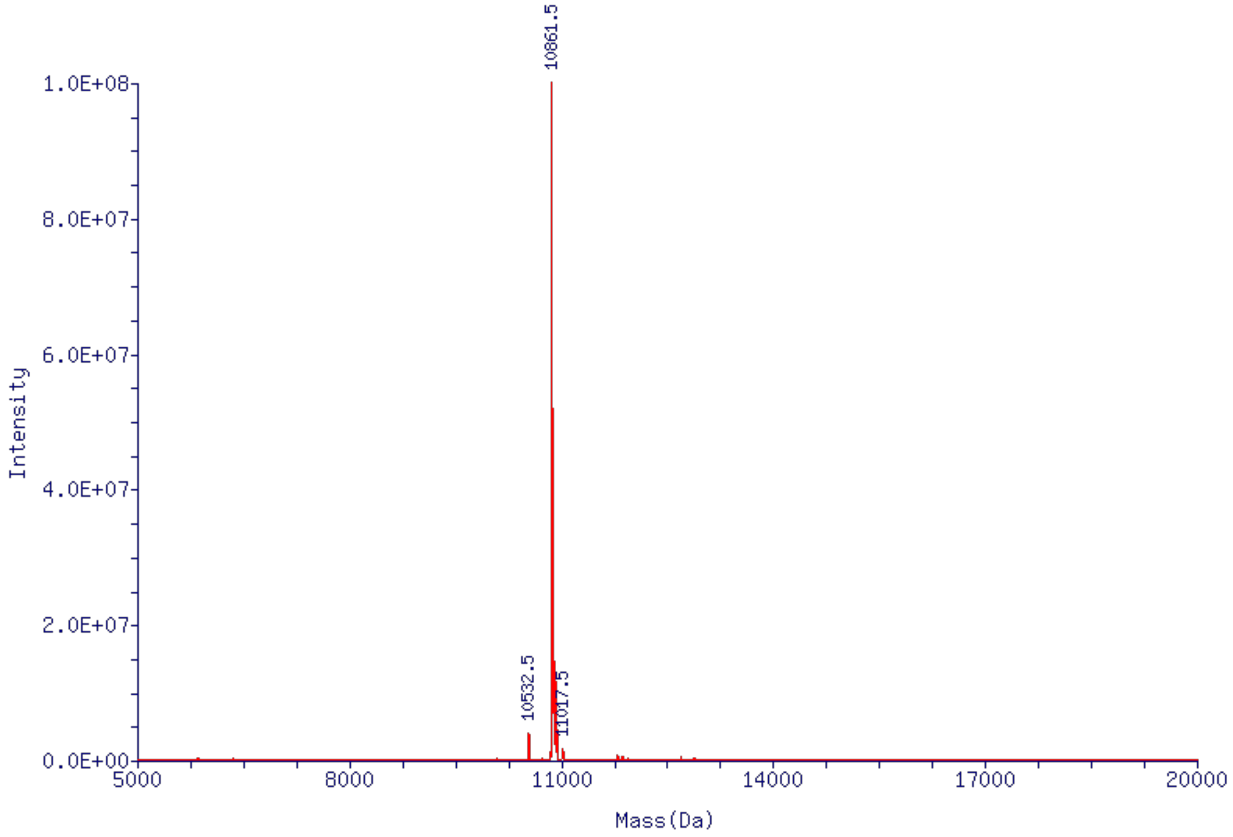


**Figure S11**. ESI-MS spectrum of BZ-5caCFWD. Mass calculated: 10860.17 Da, Mass found: 10861.5 Da.

**Figure S12**

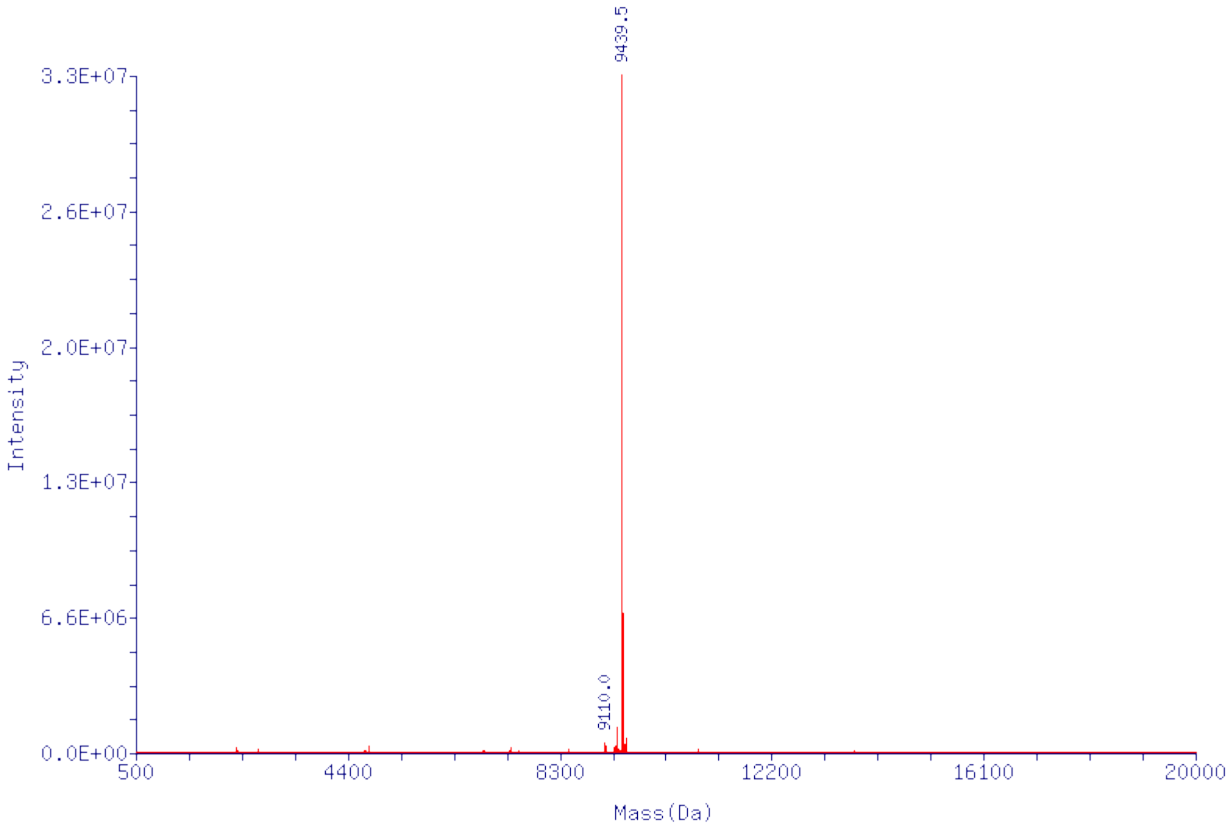


**Figure S12**. ESI-MS spectrum of AD-dU$^{F}$FWD. Mass calculated: 9439.24 Da, Mass found: 9439.5 Da.

**Figure S13**

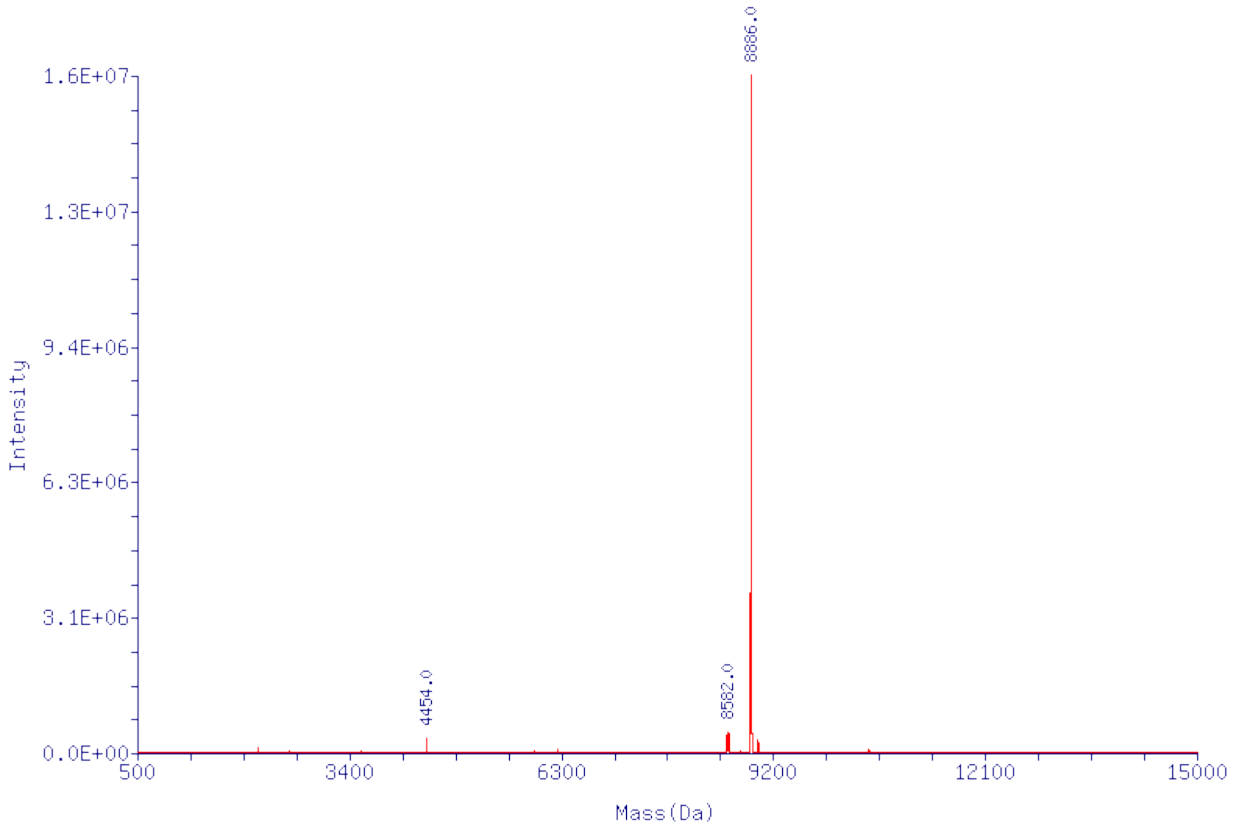


**Figure S13**. ESI-MS spectrum of AD-dT$^{F}$FWD. Mass calculated: 8886.78 Da, Mass found: 8886.0 Da.

**Figure S14**

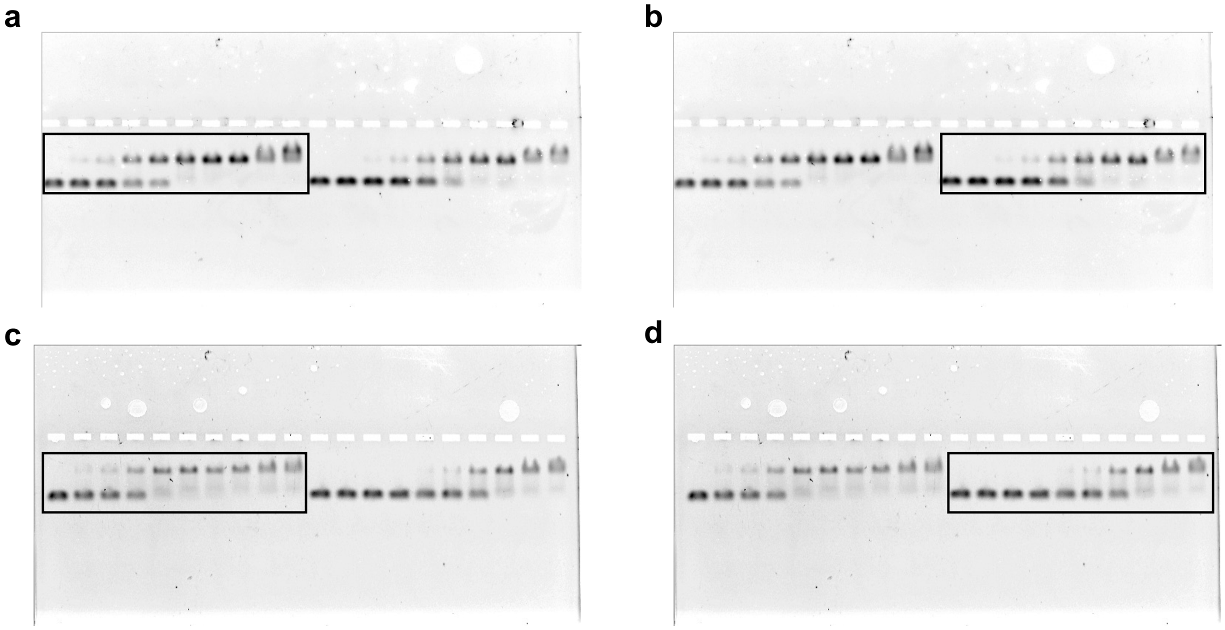


**Figure S14.** Uncropped gel images for the main text Figure 1b and 1c. (a) D1-U$^F$. (b) H1-U$^F$. (c) D2-U$^F$. (d) H2-U$^F$. Box regions indicate the cropped image shown in Figure 1b and 1c.

**Figure S15**

a b

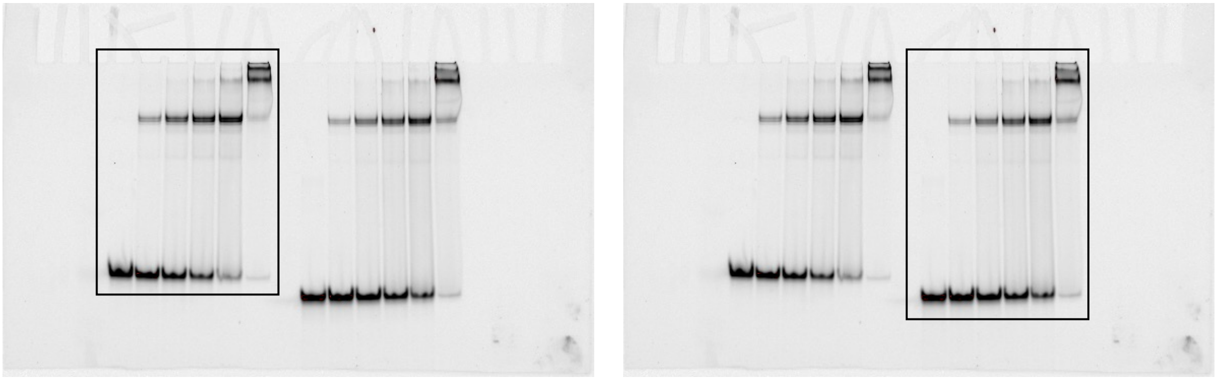

**Figure S15.** Uncropped gel images for Figure S1. (a) H1-U$^F$. (b) H2-U$^F$. Box regions indicate the cropped image shown in Figure S1.

**Figure S16**

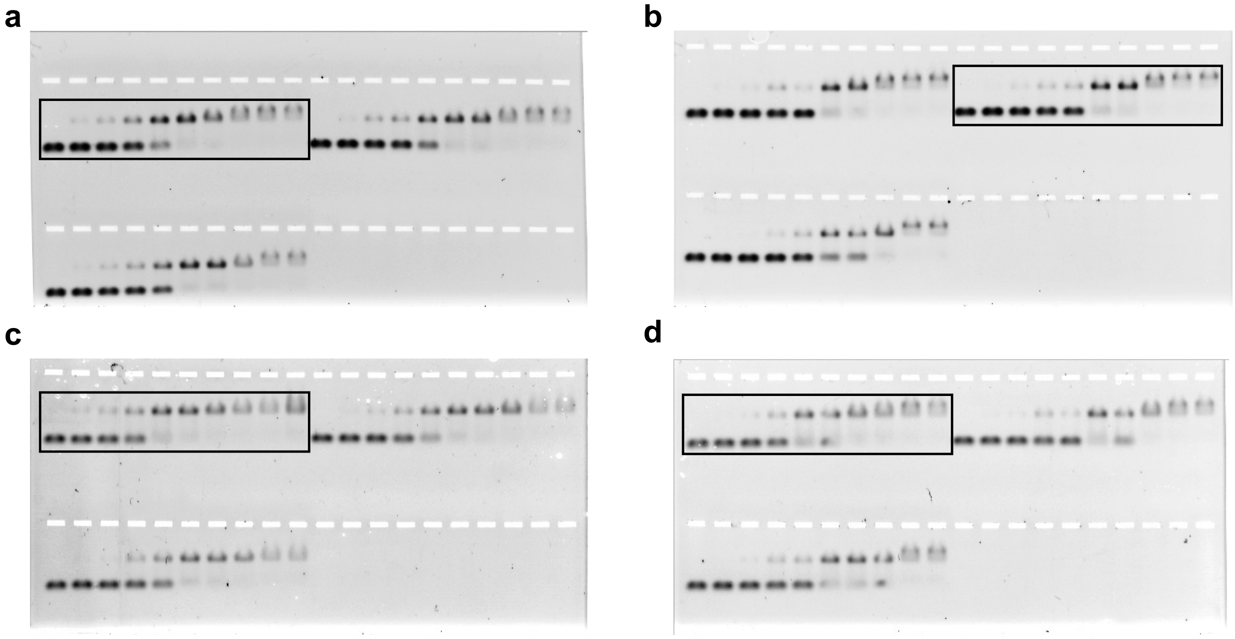

**Figure S16**. Uncropped gel images for Figure S2. (a) D1-C. (b) H1-C. (c) D2-C. (d) H2-C. Box regions indicate the cropped image shown in Figure S2.

**Figure S17**

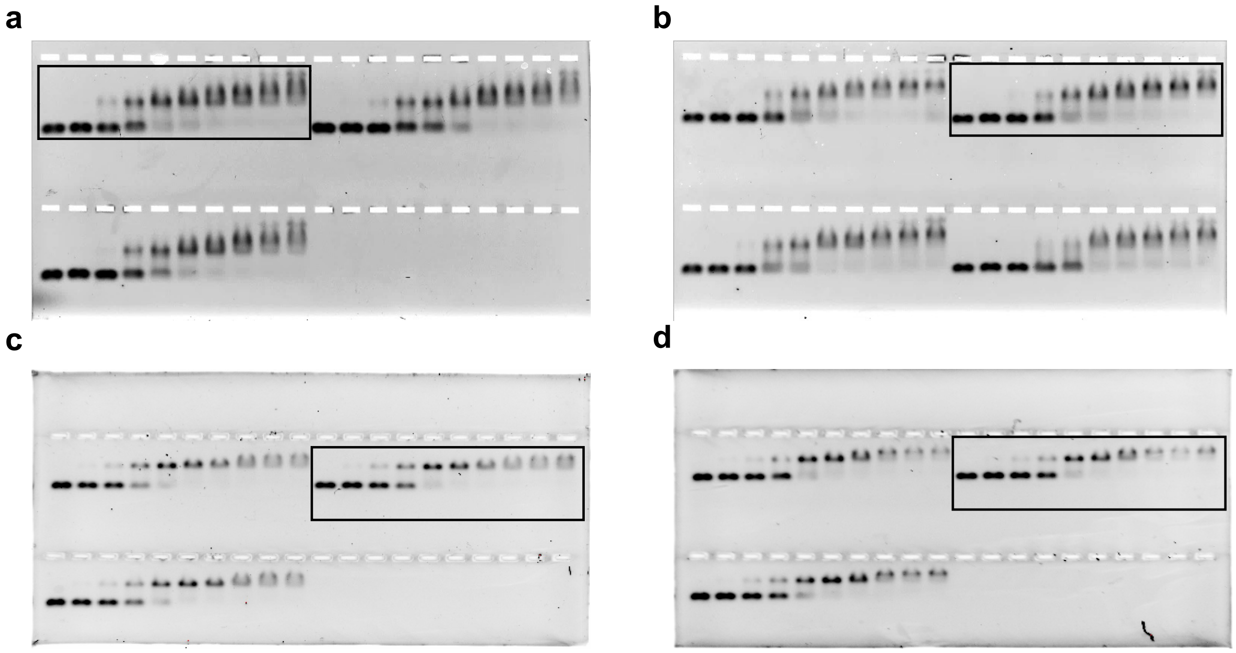


**Figure S17**. Uncropped gel images for Figure S4. (a) D1-5fC. (b) H1-5fC. (c) D2-5caC. (d) H2-5caC. Box regions indicate the cropped image shown in Figure S4.

### Figure S18

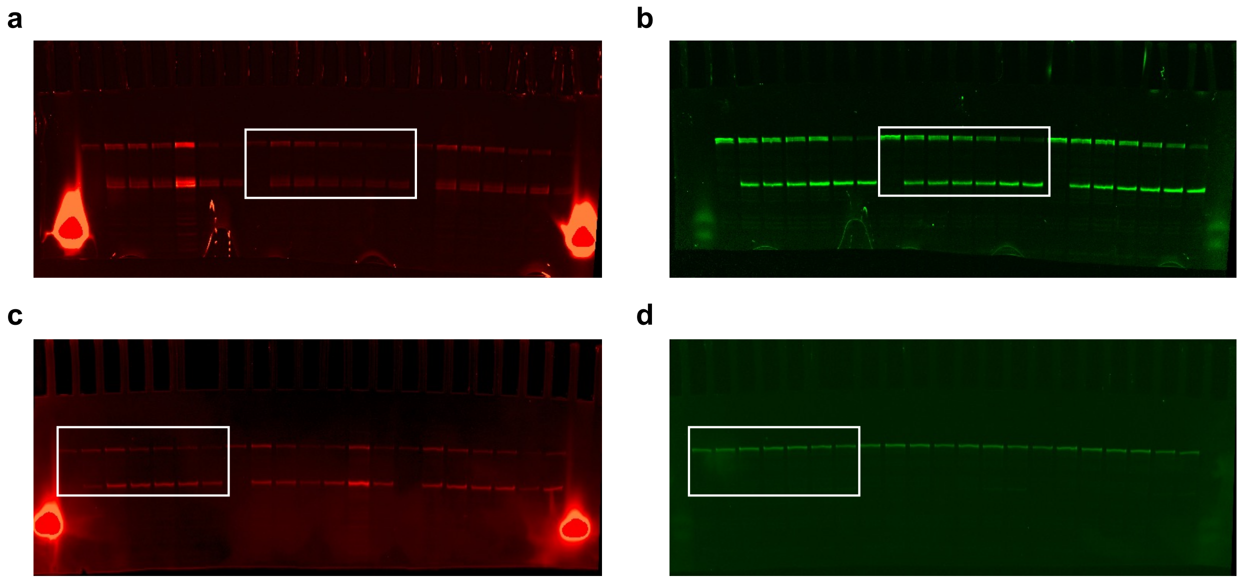


**Figure S18**. Uncropped gel images for Figure S7. (a) symGU (Cy5). (b) symGU (Cy3). (c) symGU-R (Cy5). (d) symGU-R (Cy3). Box regions indicate the cropped image shown in Figure S7.

**Figure S19**

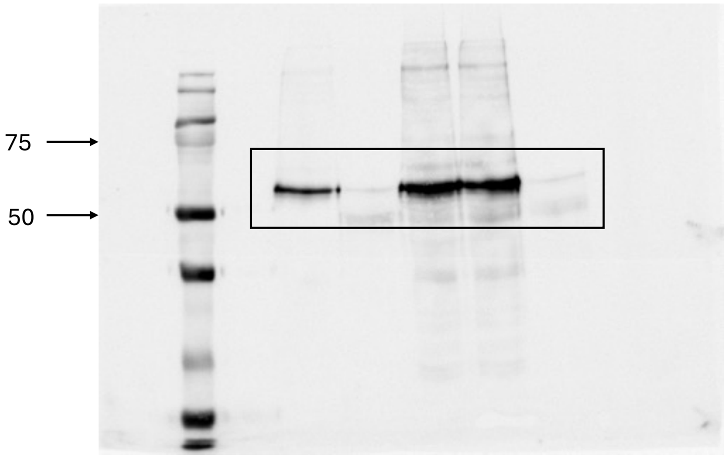


**Figure S19**. Uncropped gel image for Figure 7e. Frame indicates the cropped regions shown in Figure 7e. The calculated molecular weight of TDG is 46 kDa.

### S2. Supplementary Tables

**Table S1.** Names and sequences of all oligonucleotides used in this work. U = deoxyuridine; T = deoxythymidine; $U^F$ = 2′-deoxy-2′-fluoroarabinouridine; $T^F$ = 2′-deoxy-2′-fluoroarabinothymidine; 5fC = 5-Formyl-2'-deoxycytidine, 5caC = 5-Carboxy-2'-deoxycytidine; /FAM/ = fluorescein /Cy5/ = sulfo-Cyanine 5 dye; /Cy3/ = sulfo-Cyanine 3 dye.

| Substrate Name | Strand Name | Sequence Identity |
|---|---|---|
| D1-U | BZ-dUFWD | /FAM/TGAGGATGTATATATCTGA**U**GCGCCGGTGGAGC |
| | BZ-REV | GCTCCACCGGCGCGTACGATATATACATCCTCA |
| H1-U | BZ-dUFWD | /FAM/TGAGGATGTATATATCTGA**U**GCGCCGGTGGAGC |
| | BZ-rREV | GCUCCACCGGCGCGUACGAUAUAUACAUCCUCA |
| D1-T | BZ-dTFWD | /FAM/TGAGGATGTATATATCTGA**T**GCGCCGGTGGAGC |
| | BZ-REV | GCTCCACCGGCGCGTACGATATATACATCCTCA |
| H1-T | BZ-dTFWD | /FAM/TGAGGATGTATATATCTGA**T**GCGCCGGTGGAGC |
| | BZ-rREV | GCUCCACCGGCGCGUACGAUAUAUACAUCCUCA |
| D1-C | BZ-dCFWD | /FAM/TGAGGATGTATATATCTGA**C**GCGCCGGTGGAGC |
| | BZ-REV | GCTCCACCGGCGCGTACGATATATACATCCTCA |
| H1-C | BZ-dCFWD | /FAM/TGAGGATGTATATATCTGA**C**GCGCCGGTGGAGC |
| | BZ-rREV | GCUCCACCGGCGCGUACGAUAUAUACAUCCUCA |
| D1-5fC | BZ-5fCFWD | /FAM/TGAGGATGTATATATCTGA/**5fC**/GCGCCGGTGGAGC |
| | BZ-REV | GCTCCACCGGCGCGTACGATATATACATCCTCA |
| H1-5fC | BZ-5fCFWD | /FAM/TGAGGATGTATATATCTGA/**5fC**/GCGCCGGTGGAGC |
| | BZ-rREV | GCUCCACCGGCGCGUACGAUAUAUACAUCCUCA |
| D1-5caC | BZ-5caCFWD | /FAM/TGAGGATGTATATATCTGA/**5caC**/GCGCCGGTGGAGC |
| | BZ-REV | GCTCCACCGGCGCGTACGATATATACATCCTCA |
| H1-5caC | BZ-5caCFWD | /FAM/TGAGGATGTATATATCTGA/**5caC**/GCGCCGGTGGAGC |
| | BZ-rREV | GCUCCACCGGCGCGUACGAUAUAUACAUCCUCA |

| | | |
|---|---|---|
| D1-U$^F$ | BZ-dU$^F$FWD | /FAM/TGAGGATGTATATATCTGA/$\mathbf{U^F}$/GCGCCGGTGGAGC |
| | BZ-REV | GCTCCACCGGCGCGTACGATATATACATCCTCA |
| H1-U$^F$ | BZ-dU$^F$FWD | /FAM/TGAGGATGTATATATCTGA/$\mathbf{U^F}$/GCGCCGGTGGAGC |
| | BZ-rREV | GCUCCACCGGCGCGUACGAUAUAUACAUCCUCA |
| D2-U | AD-dUFWD | /FAM/TGTGTCACCACTGCTCA**U**GTACAGAGCTG |
| | AD-REV | CAGCTCTGTACGTGAGCAGTGGTGACAC |
| H2-U | AD-dUFWD | /FAM/TGTGTCACCACTGCTCA**U**GTACAGAGCTG |
| | AD-rREV | CAGCUCUGUACGUGAGCAGUGGUGACAC |
| D2-T | AD-dTFWD | /FAM/TGTGTCACCACTGCTCA**T**GTACAGAGCTG |
| | AD-REV | CAGCTCTGTACGTGAGCAGTGGTGACAC |
| H2-T | AD-dTFWD | /FAM/TGTGTCACCACTGCTCA**T**GTACAGAGCTG |
| | AD-rREV | CAGCUCUGUACGUGAGCAGUGGUGACAC |
| D2-C | AD-dCFWD | /FAM/TGTGTCACCACTGCTCA**C**GTACAGAGCTG |
| | AD-REV | CAGCTCTGTACGTGAGCAGTGGTGACAC |
| H2-C | AD-dCFWD | /FAM/TGTGTCACCACTGCTCA**C**GTACAGAGCTG |
| | AD-rREV | CAGCUCUGUACGUGAGCAGUGGUGACAC |
| D2-U$^F$ | AD-dU$^F$FWD | /FAM/TGTGTCACCACTGCTCA/$\mathbf{U^F}$/GTACAGAGCTG |
| | AD-REV | CAGCTCTGTACGTGAGCAGTGGTGACAC |
| D2-T$^F$ | AD-dT$^F$FWD | TGTGTCACCACTGCTCA/$\mathbf{T^F}$/GTACAGAGCTG |
| | AD-REV | CAGCTCTGTACGTGAGCAGTGGTGACAC |
| TCF21-DNA | BZ-RL1 | /Cy5/TCCTAGTGTCCACCAAATTCCTCAGCGCT**U**GCTCACCCTCCTCTACGGCCACGACTCTGGGAGTG |
| | BZ-RL2 | /Cy3/CACTCCCAGAGTCGTGGCCGTAGAGGAGGGTGAGCGAGCGCTGAGGAATTTGGTGGACACTAGGA |
| TCF21-hybrid | BZ-RL1 | /Cy5/TCCTAGTGTCCACCAAATTCCTCAGCGCT**U**GCTCACCCTCCTCTACGGCCACGACTCTGGGAGTG |
| | BZ-rRL(RNA) | GAGGGUGAGCGAGCG |
| | BZ-RL21 | CTGAGGAATTTGGTGGACACTAGGA |

| | | |
|---|---|---|
| | BZ-RL22 | CACTCCCAGAGTCGTGGCCGTAGAG |
| TCF21-loop | BZ-RL1 | /Cy5/TCCTAGTGTCCACCAAATTCCTCAGCGCT**U**GCTCACCCTCCTCTACGGCCACGACTCTGGGAGTG |
| | BZ-RL2con | CACTCCCAGAGTCGTGGCCGTAGAGTCTTAAAGTATGATACTGAGGAATTTGGTGGACACTAGGA |
| | BZ-rRL(RNA) | GAGGGUGAGCGAGCG |
| TCF21-flap | BZ-RL1 | /Cy5/TCCTAGTGTCCACCAAATTCCTCAGCGCT**U**GCTCACCCTCCTCTACGGCCACGACTCTGGGAGTG |
| | BZ-RL21F | TCTTAAAGTATGATACTGAGGAATTTGGTGGACACTAGGA |
| | BZ-RL22 | CACTCCCAGAGTCGTGGCCGTAGAG |
| | BZ-rRL(RNA) | GAGGGUGAGCGAGCG |
| TCF21-loopEX | BZ-RL1 | /Cy5/TCCTAGTGTCCACCAAATTCCTCAGCGCT**U**GCTCACCCTCCTCTACGGCCACGACTCTGGGAGTG |
| | BZ-RL2EX | CACTCCCAGAGTCGTGGCCGTAGAGTCTTAAAGTATGATATCTTAAAGCACTATACTGAGGAATTTGGTGGACACTAGGA |
| | BZ-rRL(RNA) | GAGGGUGAGCGAGCG |
| symGU | BZ-NRL1 | /Cy5/ATTGCCCTCGAGGTACCATGGATCCGATGT**U**GACCTCAAACCTAGACGAATTCCGTAGAC |
| | BZ-NRL2 | /Cy3/GTCTACGGAATTCGTCTAGGTTTGAGGT**U**GACATCGGATCCATGGTACCTCGAGGGCAAT |
| symGU-R | BZ-NRL1 | /Cy5/ATTGCCCTCGAGGTACCATGGATCCGATGT**U**GACCTCAAACCTAGACGAATTCCGTAGAC |
| | BZ-NRL2con | /Cy3/TAACGGGAGCTCCATGGTACCTTCTTGCATGTCCC**U**ATTTGGATCTGCTTAAGGCATCTG |
| | BZ-rNRL (RNA) | GAGGUCGACAUCGGA |
